# G-quadruplex formation in long non-coding RNAs dysregulated in colorectal cancer

**DOI:** 10.1101/2024.07.04.602106

**Authors:** Shubham Sharma, Chinmayee Shukla, Jérémie Mitteaux, Angélique Pipier, Marc Pirrotta, Marie-José Penouilh, David Monchaud, Bhaskar Datta

## Abstract

Non-coding RNAs (ncRNAs) in human cells do not lead to protein synthesis and constitute a substantial portion of the transcriptome. Human long non-coding RNAs (lncRNAs) orchestrate critical cellular functions influencing development, differentiation, and metabolism. Dysregulation of lncRNAs has been correlated with several pathological conditions such as neurodegenerative and autoimmune disorders, diabetes, and cancer. Recent reports have suggested the involvement of G4s in lncRNAs to regulate colorectal cancer (CRC) carcinogenesis. In this study, we investigate the occurrence and distribution of G4s in the *LINC01589*, *MELTF-AS1,* and *UXT-AS1* lncRNAs, which have been reported to be dysregulated in CRC. Using a combination of *in silico* tools and *in vitro* biophysical techniques, we show that these lncRNAs form stable, parallel, and intramolecular G4s. Furthermore, we establish the formation of G4s within these lncRNAs in CRC using cell-based assays, including RNA G4-Immuno-FISH and G4RP-RT-qPCR. This is the first systematic study of G4s in lncRNAs dysregulated in CRC, and our findings highlight the diagnostic and therapeutic potential of G4s in CRC.

## Introduction

Long non-coding RNAs (lncRNAs) represent an important class of ncRNAs within the human transcriptome, encompassing pivotal functions in both normal cellular physiology and disease pathogenesis.^1,2^ These RNAs are characterized by their length (> 200 nucleotides), translationally silent nature, and tissue-specific expression patterns. Their unique and intricate three-dimensional structures, in conjunction with finely orchestrated spatiotemporal expression patterns, make lncRNAs key players in the regulation of cell growth, differentiation, and developmental processes, acting at both genetic and epigenetic levels.^3–6^ These central regulatory roles also explain why lncRNAs are often dysregulated in various pathologies, including cancers, thus holding a privileged place as putative targets for both diagnostic and therapeutic interventions.^7–10^

Among the structural motifs lncRNAs are replete with, G-quadruplexes (G4s) rank high.^11–13^ G4s are thermodynamically stable secondary structures that fold from guanine (G)-rich sequences owing to the ability of Gs to self-associate to form G-quartets, which then self-stack to form the G4 (Figure 1).^14–17^ The ncRNA G4s are involved in several ncRNA-related processes, from the docking of RNA binding proteins (RBPs) to miRNA maturation, from the control of the ncRNA stability to that of its cellular localization. Significant efforts are currently being invested to accurately characterize the cellular G4-modulators, chaperones, and helicases.^11–13,18^

**Figure 1.**
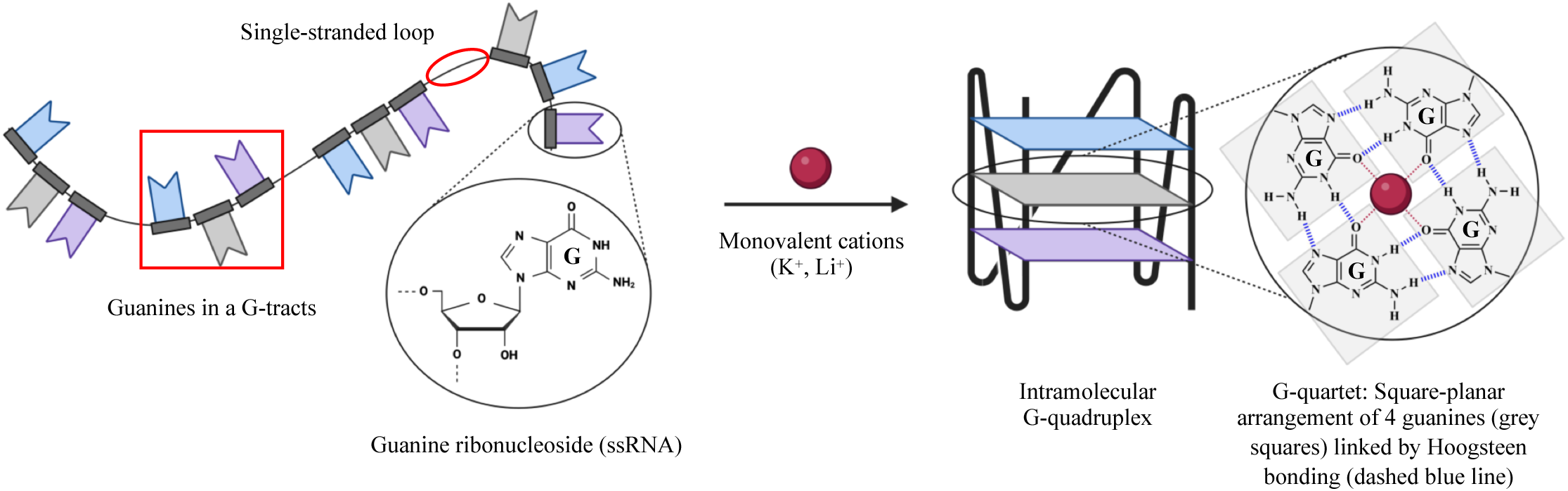
G-quadruplex (G4) structure formation. G4-formation involves Hoogsteen bonding between guanines, resulting in planar G-quartets. G-quartets stack on top of one another and enclose suitable coordination sites for monovalent cations. A number of contiguous guanines and intermittent loop sequences contribute to the overall G4 structure.

*In silico* transcriptome-wide analyses first shed light on very high propensity of G4-formation in lncRNAs.^19^ The demonstration of their existence in cells was challenging, owing to their dynamic regulation by chaperones and helicases, which make G4-formation transient.^11,18^ Recent findings within *NEAT1* and *MALAT1* lncRNAs substantiated the roles that G4s may play in the ncRNA biology.^20–22^ However, further efforts are required to understand the exact contribution of G4s in the functions of the cognate lncRNAs, and to identify their interacting partners.

We decided to focus on lncRNAs found to be dysregulated in colorectal cancer (CRC) as it is the second-highest cause of mortalities related to cancer worldwide, with > 900,000 deaths each year.^23^ While recent improvements in therapies and surgery have resulted in a 5-year survival rate of ∼ 64%, the molecular basis of CRC progression remains unclear at present.^24,25^ LncRNAs play a prominent role in CRC progression, and their over-expression, down-regulation, or mutations have been leveraged towards CRC diagnosis and prognosis, and for explaining tumorigenesis or metastasis.^26,27^ By virtue of their secondary structures, lncRNAs can interact with DNAs, RNAs, and proteins, serving as oncogenes or tumor-suppressor genes by modulating cell proliferation, apoptosis, epithelial-mesenchymal transition (EMT), angiogenesis, and metastasis in CRC.^28^ The tissue-specific expression of lncRNAs and their traceability in various bodily fluids marks them as potent biomarkers.^29,30^ In spite of these appealing characteristics, the dysregulation of lncRNAs in CRC is less explored compared to other RNAs, and the questions pertaining to their mechanism of action and effects on CRC pathogenesis remain unanswered.^26,27,31^

Recent reports suggest the involvement of G4s in CRC-dysregulated lncRNAs (*GSEC*, *REG1CP*, and *LUCAT1*) in the regulation of CRC pathogenesis.^32–34^ These studies indicate that G4s within these lncRNAs may provide a hub for binding various regulatory entities associated with CRC progression. Recently, lncRNAs, including *LINC01589*, *MELTF-AS1*, and *UXT-AS1*, have been identified as dysregulated in CRC, with implications in cell proliferation, migration, and apoptosis in CRC.^35–37^ However, the G4-formation in such lncRNAs and the broader impact of their G4s on CRC modulation have yet to be explored, which formed the basis of our current investigation. In this work, we follow a novel *in silico* pipeline to identify G4s in lncRNAs dysregulated in CRC and shortlist *LINC01589*, *MELTF-AS1*, and *UXT-AS1* lncRNAs for further investigation. The expression of these lncRNAs in cancers other than CRC indicates their potential role in diverse cancers. We show that G4s within these lncRNAs are stable and fold into parallel and intramolecular conformation. The number of constitutive G-quartets in the lncRNAs studied plays a significant role in stabilizing their respective G4s. We demonstrate G4-folding by the lncRNAs in CRC using a combination of cell-based assays, including immunofluorescence and fluorescence *in situ* hybridization (RNA G4-Immuno-FISH), and G4-RNA-specific precipitation (G4RP-RT-qPCR).^18,38–45^ The cytoplasmic prevalence of these lncRNAs and their moderate G4-formation in both cytoplasm and nucleus suggest the dynamic or transient character of these G4s.^18^ This is the first report that systematically characterizes the G4-formation in CRC-dysregulated *LINC01589*, *MELTF-AS1*, and *UXT-AS1* lncRNAs *in vitro* and *in cella*. The identified G4s can be exploited as anchors to isolate lncRNAs and study their roles and implications in various pathologies.

## Materials and Methods

### Oligonucleotides

DNA oligonucleotides corresponding to T7 RNA promoter (sense strand), cDNA antisense templates for *in vitro* transcription (IVT), 5ʹ Texas Red-labelled primer for RT stop assay, and competitor DNA oligonucleotides were meticulously designed using SnapGene software (Dotmatics, www.snapgene.com, USA), and procured from Sigma-Aldrich Chemicals Pvt. Ltd., Bangalore, India. RNA oligonucleotides corresponding to the Putative G-quadruplex-forming Sequences (PQS) of selected lncRNAs along with the FAM-21-TAMRA (F21T) DNA oligonucleotide were sourced from Kaneka Eurogentec S.A., Belgium. DNA primers for Reverse Transcription-quantitative Polymerase Chain Reaction (RT-qPCR) and 5ʹ 6-FAM-labelled DNA probes for RNA G4 Immunofluorescence and Fluorescence *in situ* hybridization (RNA G4-Immuno-FISH) were designed using Primer-BLAST (NCBI, USA) and acquired from Kaneka Eurogentec S.A., Belgium, and Eurofins Genomics India, Bangalore, India, respectively.^46^ The oligonucleotides were reconstituted in the nuclease-free water to a final concentration of 100 – 500 µM and stored at −20 °C until further use. Details regarding the sequences of the DNA and RNA oligonucleotide mentioned above are accessible in Table S2-4.

### Reagents

The HiScribe^™^ T7 High Yield RNA Synthesis Kit (Catalog no. E2040S), DNase I (Rnase-free) (Catalog no. M0303L), Monarch^®^ RNA Cleanup Kit (Catalog no. T2050L), Rnase Inhibitor, Murine (Catalog no. M0314L), and Ribonucleoside Vanadyl Complex (Catalog no. S1402S) were procured from New England Biolabs Pte. Ltd., Singapore. Thioflavin T (Catalog no. T3516), RNase A (Catalog no. 10109169001), TMPyP4 (Catalog no. 613560), Bovine Serum Albumin (Catalog no. A9418), and Ribonucleic acid, transfer from baker’s yeast (S. cerevisiae) (Catalog no. R8759) were obtained from Sigma-Aldrich Chemicals Pvt. Ltd., Bangalore, India. Biotin (Catalog no. B4501) was acquired from Sigma Aldrich Chimie S.a.r.l, France. Deoxynucleotide Triphosphates (dNTPs) Mix (Catalog no. U1511) and M-MLV Reverse Transcriptase (RNase H Minus) (Catalog no. M5301) were procured from Promega Biotech India Pvt. Ltd, New Delhi, India. Streptavidin MagneSphere^®^ Paramagnetic Particles (Catalog no. Z5481) were obtained from Promega France, France. TRIzol™ Reagent (Catalog no. 15596018), SuperScript™ III Reverse Transcriptase (Catalog no. 18080093), RNaseOUT™ Recombinant Ribonuclease Inhibitor (Catalog no. 10777019), Random Hexamers (Catalog no. N8080127), and Oligo(dT)_20_ Primer (Catalog no. 18418020) were acquired from Thermo-Fischer Scientific Inc., Fisher Scientific SAS, France. iTaq Universal SYBR Green Supermix (Catalog no. 1725121) was procured from Bio-Rad, France. Dextran Sulphate Sodium Salt 500 (DSS 500) ex. Leuconostoc Sp. (Catalog no. 99629) was obtained from Sisco Research Laboratories Pvt. Ltd., Mumbai, India. Anti-DNA G-quadruplex structures Antibody, clone BG4 (Catalog no. MABE917), DYKDDDDK Tag (D6W5B) Rabbit mAb (Catalog no. 14793), and Goat anti-Rabbit IgG (H+L) Cross-Adsorbed Secondary Antibody, and Alexa Fluor™ 568 (Catalog no. A-11011) were procured from Sigma-Aldrich Chemicals Pvt. Ltd., Bangalore, India, Cell Signaling Technology, Inc., Research Instruments Pte. Ltd., Singapore, and Thermo Fischer Scientific India Pvt. Ltd., Mumbai, India, respectively. Dulbecco’s Modified Eagle Medium (Catalog no. L0104) and Fetal bovine serum (FBS) (Catalog no. S1810) were obtained from Dominique DUTSCHER SAS, France and Thermo Fischer Scientific India Pvt. Ltd., Mumbai, India. RPMI 1640 Medium (Catalog no. 21870076), Leibovitz’s L-15 Medium (Catalog no. 41300039), GlutaMAX™ (Catalog no. 35050061), L-Glutamine (Catalog no. 25030149), and Penicillin-Streptomycin (Catalog no. 15140122) were acquired from Thermo-Fischer Scientific Inc., Fisher Scientific SAS, France, and Thermo Fischer Scientific India Pvt. Ltd., Mumbai, India. Naptho-TASQ and BioCyTASQ were kindly gifted by the Monchaud Lab (GATTACA), UMR CNRS 6302, ICMUB, France. All the reagents were prepared following the manufacturer’s protocol and stored in the recommended conditions.

### Cell lines

Colorectal cancer cell lines, including COLO205, COLO320DM, HCT-15, HT-29, HCT-116, and SW620, were procured from the National Cell Repository for cell lines at the National Centre for Cell Science (NCCS), Pune, India. Cell lines from breast cancer [HCC-1954 (CRL-2338), MDA-MB-231 (HTB-26), MCF7 (HTB-22)], prostate cancer [PC3 (CRL-1435)] and cervical cancer [HeLa (CCL-2)] were acquired from the American Type Culture Collection (ATCC), USA. HT-29, HCT-116, HeLa, and MCF7 were cultured in Dulbecco’s Modified Eagle Medium (DMEM), while COLO205, COLO320DM, HCT-15, HCC-1954, MDA-MB-231, and PC3 were cultured in RPMI 1640 Medium. SW620 was cultured in Leibovitz’s L-15 Medium. All cell lines were supplemented with 10% (v/v) Fetal Bovine Serum and 1% (v/v) Penicillin-Streptomycin. Additionally, cell lines cultured in DMEM and RPMI 1640 medium were supplemented with 1% (v/v) GlutaMAX™ or L-Glutamine. All the cell lines were maintained at 37 °C in the presence of 5% CO_2_, except SW620, which was maintained in the absence of 5% CO_2_.

### *In silico* identification of Putative Quadruplex-forming Sequences (PQS) in CRC-dysregulated lncRNAs

The Lnc2Cancer 3.0 database was used to obtain the list of all the lncRNAs dysregulated in CRC.^47^ Subsequently, the FASTA sequences of all these lncRNAs and their transcript variants were retrieved from the NCBI Nucleotide database.^48^ The QGRS mapper tool was used to identify PQS within the sequences of the selected lncRNAs and their transcript variants. The parameters used for QGRS mapper were, max length: 45; min G-group: 2; loop size: 0 to 36.^49^ The G4Hunter tool was used with the following parameters: window size: 45; threshold: 0.9.^50^ The highest-scoring PQS within the selected lncRNAs were chosen for *in vitro* investigation.

### RNA isolation and Reverse Transcription-quantitative Polymerase Chain Reaction (RT-qPCR)

RNA isolation from the mentioned cell lines was performed using the conventional TRIzol method.^51^ Following the RNA purification and DNase treatment of the purified RNAs, reverse transcription was carried out using the SuperScript™ III Reverse Transcriptase, Random Hexamers, and Oligo(dT)_20_ Primer, following the manufacturer’s protocol. The resulting cDNAs were used to perform the quantitative Polymerase Chain Reaction (qPCR) with the iTaq Universal SYBR Green Supermix, as per the manufacturer’s protocol. The qPCR was carried out in the presence of primer pairs specific for the selected lncRNAs and housekeeping genes (GAPDH and β-Actin) using the Mx3005P qPCR System (Table S2). The data were recorded in triplicates across two independent studies to determine lncRNA expression levels in CRC (HT-29, HCT-116), breast cancer (HCC-1954, MDA-MB-231, MCF7), prostate cancer (PC3), and cervical cancer (HeLa) cell lines. The mean relative lncRNA expression (2^Δ1Ct^) with respect to GAPDH were plotted with standard error of mean against different cancer cell lines using GraphPad Software. For assessing lncRNA expression levels in different CRC cell lines (HT-29, HCT-116, HCT-15, COLO205, COLO320DM, and SW620), the data were recorded in triplicate as part of an independent study. The mean relative lncRNA expression (2^Δ1Ct^) with respect to β-Actin were plotted with standard deviation against different CRC cell lines using GraphPad Software.

### Preparation of wild-type and deletion mutants of lncRNAs

The templates for IVT were designed by sequentially incorporating the antisense sequences of transcription initiator, the cDNA of wild-type (WT) or deletion mutant (Δ) of respective lncRNA PQS, ten extra bases, and 20-nt long primer binding site toward the 5ʹ-end of the antisense sequence of the T7 RNA promoter. The deletion mutants of lncRNA PQS were devoid of singular G-tracts. To minimize heterogeneity and prevent the incorporation of additional G-tract to the existing PQS, the conventional transcription initiation sequence CCCTTT was replaced with CGCTTT in the cDNA antisense templates.^52^ Equimolar concentrations of the resuspended T7 RNA promoter (sense strand) oligonucleotide and cDNA antisense template for wild-type and deletion mutants of respective lncRNA PQS were mixed in an annealing buffer: 10 mM Tris-HCl (pH 7.5), 50 mM NaCl and 1 mM EDTA (pH 8.0), and annealed by heating at 95 °C, followed by gradual cooling to room temperature (Table S3). The absorbance measurements at 260 nm, A_260/280_, and A_260/230_ values were obtained using a NanoDrop™ 2000c spectrophotometer (Thermo Fischer Scientific India Pvt. Ltd., Mumbai, India), to determine the concentration and purity of the annealed oligonucleotides. The annealed oligonucleotides were then used to perform IVT with the HiScribe™ T7 High Yield RNA Synthesis Kit, following the manufacturer’s protocol. The transcribed RNAs underwent DNase I (RNase-free) treatment for degradation of the DNA oligonucleotides and were subsequently purified using Monarch® RNA Cleanup Kit and reconstituted in nuclease-free water. The purified RNAs were supplemented with RNase inhibitor (Murine), and their concentration and purity were assessed using a NanoDrop™ 2000c spectrophotometer. The purified IVT RNAs were stored at −80 °C until further use. The IVT RNAs (2 µM) were also run on a 15% native polyacrylamide gel electrophoresis (PAGE) to check the RNA integrity (data not shown). The IVT RNAs (5 µM) were subjected to heating at 95 °C for 5 minutes in the presence of a folding buffer: 10 mM Tris-HCl (pH 7.5) and 0.1 mM EDTA (pH 8.0), supplemented with or without specific monovalent cations (KCl – K^+^ or LiCl – Li^+^). The RNAs were then gradually cooled to room temperature to facilitate the formation of the G4s. The synthetic RNAs were reconstituted in the nuclease-free water to a final concentration of 500 µM and stored at −20 °C until further use (Table S3). A procedure analogous to that used for IVT RNAs was followed to fold the synthetic RNAs (1 µM) into G4s. While the folded synthetic RNA oligonucleotides were used for N-TASQ fluorescence enhancement assay, FRET-MC, and HILIC-ESI-MS, the folded IVT RNAs were used for other *in vitro* experiments.

### Circular Dichroism Spectroscopy

Circular dichroism (CD) spectra spanning from 220 nm to 320 nm were recorded for the wild-type RNAs (1 or 5 µM) folded in the presence or absence of additional 10 or 100 mM KCl or 100 mM LiCl, on a J-815 CD Spectropolarimeter (JASCO Inc., USA), at 25 °C. The measurement parameters were set as follows, data pitch: 1 nm; sensitivity: standard; DIT: 1 sec; bandwidth: 0.50 nm; scanning speed (100 nm/ min), and accumulations: 3. After smoothening the data with the Savitzky-Golay method and 20 points of window, the mean CD (mdeg) values were plotted against the wavelength for respective RNAs using the Origin (Pro), Version 2017 (OriginLab Corporation, USA). CD-melting experiments were conducted by thermally denaturing the wild-type RNAs (1 µM) folded in the supplementation of 10 mM KCl, in the temperature range of 25 °C to 95 °C. Simultaneous recording of the CD (mdeg) values was carried out at intervals of 1 °C on a J-815 CD Spectropolarimeter, using the parameters mentioned above. The three accumulations at a wavelength corresponding to the intense maxima obtained in the CD spectra of the respective folded RNAs were obtained at each temperature. The mean CD (mdeg) values were smoothened with the Savitzky-Golay method and 20 points of window, normalized, and plotted against the temperature for respective RNAs using Origin (Pro) and GraphPad Software (Dotmatics, USA). The plot was fitted using the dose response-inhibition model of non-linear regression to determine the melting temperatures (T_m_) of the respective RNAs. Similarly, CD spectroscopy and CD-melting of the folded RNAs (1 µM) were also conducted in the presence of BioCyTASQ (1 – 5 µM), and the data were plotted and fitted in an analogous manner.

### NaphthoTASQ (N-TASQ) Fluorescence enhancement assay

Wild-type RNAs (2 µM) folded in the supplementation of 10 mM KCl were titrated with N-TASQ (1 µM), reaching a final concentration of 5 µM. After each titration, the fluorescence emission spectrum of N-TASQ was recorded from 335 nm to 600 nm, along with endpoint fluorescence emission at 393 nm, with excitation at 280 nm using a CLARIOstar^®^ Plus multimode plate reader (BMG LABTECH SARL, France).^53^ The data were recorded in duplicate as part of an independent study (due to RNA scarcity). The mean fluorescence intensities (arbitrary units) of emission spectra were plotted against the wavelength after smoothening the data with the Savitzky-Golay method and 20 points of window using Origin (Pro). The mean N-TASQ fluorescence fold-enhancements (endpoint fluorescence in the presence of RNA/ absence of RNA: F/ F_0_) were plotted with standard deviation against respective RNAs using GraphPad Software. Ordinary one-way or two-way ANOVA were employed for the statistical analyses.

### Fluorescence resonance energy transfer-melting competition (FRET-MC) assay

Wild-type RNAs (3 µM) folded in the supplementation of 10 mM KCl were mixed with the folded F21T (0.2 µM) and N-TASQ (1 µM) in CaCoK10 buffer: 10 mM lithium cacodylate buffer (pH 7.2) and 10 mM KCl. The thermal denaturation of this mixture was carried out from 25 °C to 90 °C, with the intensity of the FAM emission simultaneously recorded at intervals of 1 °C when excited at 488 nm, using a Mx3005P qPCR System (Agilent Technologies, France).^54,55^ The data were recorded in duplicates across two independent studies. The means of normalized FAM emission at maxima from F21T in the presence or absence of RNAs and N-TASQ were plotted with standard error of mean against the temperature for respective RNAs using GraphPad Software. The plot was fitted using the Boltzmann sigmoidal model of non-linear regression to determine the melting temperatures (T_m_) of the respective RNAs.

### Hydrophilic Interaction Liquid Chromatography (HILIC) coupled with Electrospray Ionization Mass Spectrometry (ESI-MS)

Wild-type RNAs (10 µM) were folded in 100 mM Ammonium Acetate solution (pH 7.0) to avoid incompatibility with the native electrospray ionization on the usage of non-volatile solutions, and minimize any pH-induced effects.^56,57^ Methanol was added to the folded RNAs to a working concentration of 20%.^58,59^ The prepared sample was introduced into the Vanquish HPLC System (Thermo-Fischer Scientific Inc., Fisher Scientific SAS, France) with Luna 3 µM HILIC column (Catalog no. 00D-4449-B0, 100 X 2 mm, Phenomenex, France) and subjected to a gradient (10/90 to 40/ 60) of mobile phase: 20 mM Ammonium Formate (pH 3.2) and 100% Acetonitrile, for 10 minutes, to generate the chromatogram. The UV-Vis detector was used at 260 nm to determine the RNA content with respect to the HILIC chromatogram. Samples were subsequently injected into the Orbitrap Exploris™ 240 ESI-MS (Thermo-Fischer Scientific Inc., Fisher Scientific SAS, France) and analyzed in negative mode using the following parameters, flow rate: 500 µl/ min; resolution: 120000; mass/charge range: 500 to 6000; scan type: full, and RF lens: 100%. The HILIC chromatogram and UV-Vis data were used in conjunction to determine the retention time(s) where the RNAs exhibited their highest relative abundance, with the most significant absorption at 260 nm. ESI-MS spectra were generated for the identified retention time(s), and the m/z peaks with high relative abundances were analyzed for their experimental mass.

### RNA G4 Immunofluorescence and Fluorescence in situ hybridization (RNA G4-Immuno-FISH)

The 5ʹ 6-FAM-labelled DNA probes of 25-nt each were meticulously designed against the RNAs using Primer-BLAST, while excluding the regions with high-scoring PQS identified by the QGRS mapper (Table S4). HT-29 cells were seeded on sterile glass coverslips placed in a 24-well tissue culture plate and cultured at 37 °C in the presence of 5% CO_2_ for 72 hours. Subsequently, the cells were fixed with 4% paraformaldehyde in 1X Phosphate-Buffered Saline (PBS) (pH 7.0) for 10 minutes, followed by the cell permeabilization with 0.1% triton X-100 in 1X PBS supplemented with 2 mM EGTA (pH 8.0) for 10 minutes. Except for the cells designated for RNase A treatment, 2mM Ribonucleoside Vanadyl Complex was added to the permeabilization solution. The cells were then subjected to DNase I (220 U) or RNase A (5 mg/ ml) treatment at 37 °C for 90 minutes. Nuclease-treated or untreated cells were blocked with 0.5% FBS in 1X PBS supplemented with 0.1% Tween 20 (PBST) at 37 °C for 60 minutes. The cells were then incubated with Anti-DNA G-quadruplex structures Antibody, clone BG4 (1:300), followed by DYKDDDDK Tag (D6W5B) Rabbit mAb (1:500), and Goat anti-Rabbit IgG (H+L) Cross-Adsorbed Secondary Antibody, Alexa Fluor™ 568 (1:1000), in the presence of 2 mM Ribonucleoside Vanadyl Complex at 37 °C for 60 minutes each.^38,39^ Following the sequential PBST washes between consecutive antibody incubations, cells were re-fixed with 4% paraformaldehyde in 1X PBS for 5 minutes and washed with 2X Saline-Sodium Citrate (SSC) buffer (pH 7.0). A hybridization mixture consisting of 10% formamide, 10% (w/v) Dextran Sulphate Sodium Salt 500, Bovine Serum Albumin (0.2 mg/ ml), Ribonucleic acid, transfer from baker’s yeast (1 mg/ml), 5ʹ 6-FAM-labelled DNA probes (140 nM), in 2X SSC buffer, was incubated at 37 °C for 10 minutes, followed by 95 °C for 10 minutes, and immediate cooling on ice for 10 minutes. Alexa Fluor™ 568-labelled cells were subsequently incubated with the hybridization mixture at 37 °C for 16 hours within a dark and humid chamber.^40–42^ Post hybridization, cells were washed with 10 % formamide in 2X SSC buffer, followed by 2X, 1X, and 4X SSC buffer at 37 °C. The cells were stained with DAPI, and coverslips were mounted onto the glass slides. The cells were imaged under the TCS SP8 Laser Scanning Confocal Microscope (Leica Microsystems-DHR Holding India Pvt. Ltd., Mumbai, India) using 405 nm, 488 nm, and 561 nm lasers and Hybrid detectors (HyD) with theoretic emission windows for DAPI, FAM, and Alexa Fluor™ 568, respectively. The images were analyzed for the number of cells per region of interest, the number of RNA foci (6-FAM-labelled), the colocalization of RNA foci with the G4 foci (Alexa Fluor™ 568-labelled), and the intensity of G4 foci colocalizing with the RNA foci. The analyses were performed for individual optical sections of these images with respect to the DAPI staining to obtain information regarding the subcellular localization of the RNA foci. An ImageJ macro was written for the analyses and is available on GitHub (https://github.com/ICMUB/FISH-lncRNA-IF-BG4). The data were recorded over 10-30 frames (>5 cells per frame) across two independent studies. The mean number of RNA foci per cell, mean fold-colocalization of the RNA foci with the G4 foci, and means of normalized intensities of G4 foci colocalizing with the RNA foci were plotted with whiskers at 10 – 90 percentile against nuclear or cytoplasmic compartment of HT-29 cells for the respective RNAs using the GraphPad Software. Ordinary one-way ANOVA was applied for the statistical analyses.

### G4-RNA-specific precipitation coupled with the Reverse Transcription-quantitative Polymerase Chain Reaction (G4RP-RT-qPCR)

The HT-29 cells were seeded in a T-175 tissue culture flask and cultured at 37 °C in the presence of 5% CO_2_ for 72 hours. The G4RP-RT-qPCR procedure was carried out as mentioned in the recent reports.^18,43–45^ Subsequently, the cells were trypsinized, fixed with 1% paraformaldehyde, and lysed using a 27G needle and syringe. The cell lysate was centrifuged to collect the supernatant of cell lysate. The resulting cell lysate (supernatant) was then incubated with biotinylated TASQ (BioCyTASQ) (100 µM) or biotin (negative control) (100 µM) in the presence of Streptavidin MagneSphere® Paramagnetic Particles (150 µg) at 4 °C for 120 minutes. After washing the BioCyTASQ or Biotin-conjugated streptavidin beads, reverse crosslinking was carried out for both the input lysate and the bound streptavidin beads at 70 °C for 120 minutes. Following this, the input lysate and the bound streptavidin beads were snap-frozen in TRIzol™ Reagent. RNA isolation from the input lysate and the bound streptavidin beads was performed using the conventional TRIzol method.^51^ Following the RNA purification and DNase treatment of the purified RNAs, RT-qPCR was carried out in the presence of primer pairs specific for the selected RNAs, as described earlier (Table S2). The data were recorded in triplicates across three independent studies. The mean fold-change in the level of respective RNAs in BioCyTASQ (G4RP) precipitated or biotin samples relative to the input (5%) cell lysate [5 * {2^(Mean^ ^Ct^ ^input− Ct G4RP or biotin)^}] was plotted with standard error of mean using GraphPad Software. Two-way ANOVA was applied for the statistical analyses.

### Statistical significance

The statistical analyses included appropriate tests, and the resulting statistical significance (*P*-values) with *P* ≤ 0.05, *P* ≤ 0.01, *P* ≤ 0.001, and *P* ≤ 0.0001 are denoted with one asterisk (*), two asterisks (**), three asterisks (***), and four asterisks (****), respectively. Non-significant *P*-values are not represented.

## Results and Discussion

### LncRNAs dysregulated in CRC harbor Putative Quadruplex-forming Sequences

To identify the Putative G-quadruplex-forming Sequences (PQS) in CRC-dysregulated lncRNAs, the list of all the lncRNAs dysregulated in CRC was obtained from Lnc2Cancer 3.0 and the corresponding FASTA sequences were retrieved from NCBI Nucleotide database.^47,48^ Out of 952 entries, only 611 entries (corresponding to 260 unique lncRNAs) had validated or reviewed RefSeq sequences. Sequences of these lncRNAs were analyzed for their putative G4-formation using QGRS mapper using the following parameters: max length: 45; min G-group: 2; loop size: 0 to 36.^49^ While most of the lncRNAs (257 out of 260 lncRNAs, 99%) screened had at least one PQS bearing two G-quartets (2G-G4), 120 (46%) and 8 (3%) lncRNAs had at least one PQS bearing three (3G-G4) or four G-quartets (4G-G4), respectively. The G4-forming potential of these lncRNAs was also assessed with G4Hunter using the following parameters: window size: 45; threshold: 0.9.^50^ Out of all the lncRNAs screened, 134 (51%), 101 (39%), and 7 (3%) lncRNAs had at least one PQS having the potential to fold into a 2G-G4, 3G-G4, and 4G-G4, respectively. The high-scoring 2G, 3G, and 4G PQS identified in CRC-dysregulated lncRNAs: *LINC01589*, *MELTF-AS1*, and *UXT-AS1*, respectively, using QGRS mapper and G4Hunter, encouraged us to investigate their formation through *in vitro* studies (Figure 2; Table S1).

**Figure 2.**
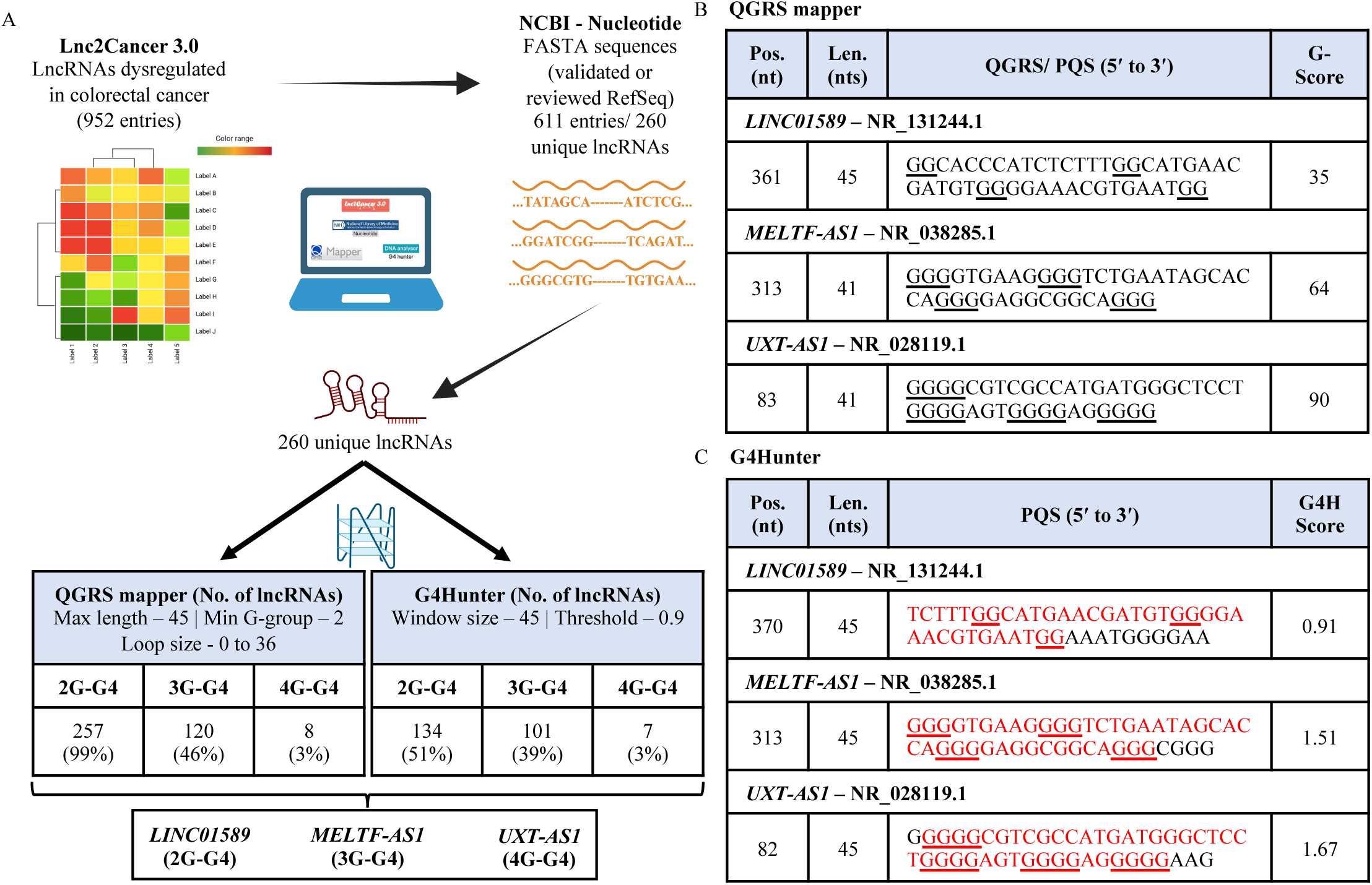
Identification of Putative Quadruplex-forming Sequences (PQS) in long non-coding RNAs (lncRNAs) dysregulated in Colorectal cancer (CRC). A) *In silico* prediction of PQS in lncRNAs dysregulated in CRC using various databases and G4-prediction tools. A list of CRC-dysregulated lncRNAs is obtained from Lnc2Cancer 3.0, and their respective FASTA sequences are obtained from NCBI nucleotide. LncRNA FASTA sequences are used in G4-prediction tools: QGRS mapper and G4Hunter, to obtain PQS. A 2G, 3G, and 4G PQS from *LINC01589*, *MELTF-AS1*, and *UXT-AS1* lncRNAs, respectively, are shortlisted for *in vitro* investigation. B-C) PQS identified from selected lncRNAs using B) QGRS mapper, and C) G4Hunter. Regions of PQS obtained from QGRS mapper overlapping with PQS from G4Hunter are marked in red.

### Expression of *LINC01589*, *MELTF-AS1*, and *UXT-AS1* lncRNAs in different cancers

We next assessed the expression levels of *LINC01589*, *MELFT-AS1*, and *UXT-AS1* lncRNAs in different cancer cell lines. To this end, we first performed the Reverse Transcription-quantitative Polymerase Chain Reaction (RT-qPCR) using RNA isolated from CRC (HT-29, HCT-116), breast cancer (HCC-1954, MDA-MB-231, MCF7), prostate cancer (PC3), and cervical cancer (HeLa) cell lines in the presence of suitable primer pairs (Table S2). The mean relative lncRNA expression with respect to GAPDH ranged from: 0.53 to 0.07 for *LINC01589* (highest: HCC-1954, lowest: MDA-MB-231 and HT-29); 1.30 to 0.12 for *MELTF-AS1* (highest: HCT-116, lowest: HT-29); 0.24 to 0.01 for *UXT-AS1* (highest: PC3, lowest: MDA-MB-231 and HCT-116) (Figure 3A). This result indicates a varied relative lncRNA expression in different cancer cell lines. It also highlights a weak to moderate relative lncRNA expression in CRC. We further investigated this by comparing the expression levels of these lncRNAs in different CRC cell lines (HT-29, HCT-116, HCT-15, COLO205, COLO320DM, and SW620) (Table S2). The mean relative lncRNA expression with respect to β-Actin ranged from: 784.11 to 46.38 for *LINC01589* (highest: SW620, lowest: HT-29); 2251.80 to 59.27 for *MELTF-AS1* (highest: COLO320DM, lowest: HT-29); 81.68 to 15.80 for *UXT-AS1* (highest: SW620, lowest: HCT-116) (Figure 3B). These findings confirmed that the selected lncRNAs were good targets for further investigations.

**Figure 3.**
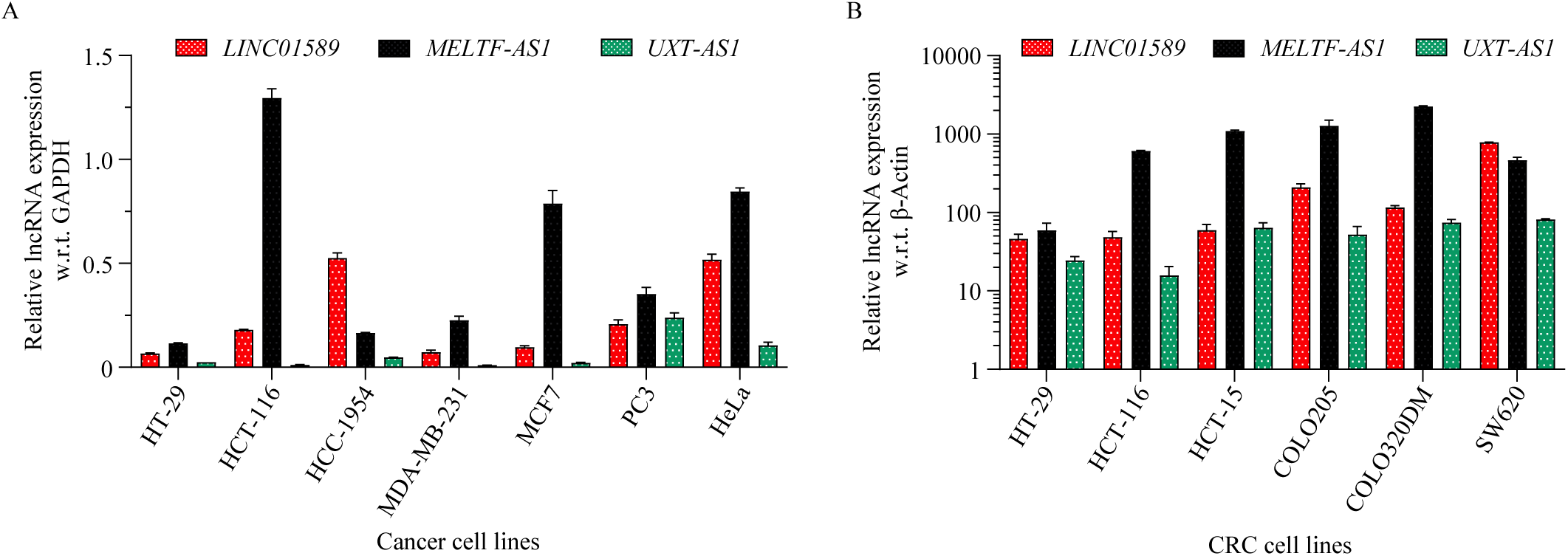
Expression of *LINC01589*, *MELTF-AS1*, and *UXT-AS1* lncRNAs in different cancers. A) Reverse Transcription-quantitative Polymerase Chain Reaction (RT-qPCR) with RNAs isolated from CRC (HT-29, HCT-116), breast cancer (HCC-1954, MDA-MB-231, MCF7), prostate cancer (PC3), and cervical cancer (HeLa) cell lines show mean relative lncRNA expression with respect to GAPDH (2^−ΔCt^) ± SEM. B) RT-qPCR with RNAs isolated from different CRC cell lines (HT-29, HCT-116, HCT-15, COLO205, COLO320DM, HT-29, and SW620) show mean relative lncRNA expression with respect to β-Actin (2^−ΔCt^) ± SD.

### *LINC01589*, *MELTF-AS1*, and *UXT-AS1* lncRNAs forms stable, parallel, and intramolecular G4s *in vitro*

To assess the *in vitro* folding of identified PQS within *LINC01589*, *MELTF-AS1*, and *UXT-AS1* lncRNAs, *in vitro* investigations were performed using *in vitro* transcription (IVT)-derived wild-type (WT) lncRNA PQS (Table S3). For simplicity, these PQS are referred to hereafter by the name of their cognate lncRNAs. CD spectroscopy is commonly used to characterize the formation and elucidate the topology of G4s in DNA and RNA.^60^ The CD signatures obtained for *LINC01589*, *MELTF-AS1*, and *UXT-AS1*, displayed the typical parallel G4 signature (maxima at *ca*. 265 and minima at *ca*. 240 nm, Figure 4A-C). CD spectroscopy was also performed on the synthetic RNAs in the presence of 10 mM KCl, and similar results were obtained (Figure S1A). These observations are in agreement with the preferred parallel topology of physiologically relevant RNA G4s.^14^ Since monovalent cations influence the G4-folding, the folding buffer for IVT-derived lncRNAs was supplemented with 100 mM KCl or LiCl.^61–63^ The addition of KCl resulted in a slightly higher 265 nm band for *MELTF-AS1* and *UXT-AS1*, but not for *LINC01589*. However, the addition of LiCl did not modify their CD signatures, which testifies to the stability of the corresponding G4s (Figure 4A-C).

**Figure 4.**
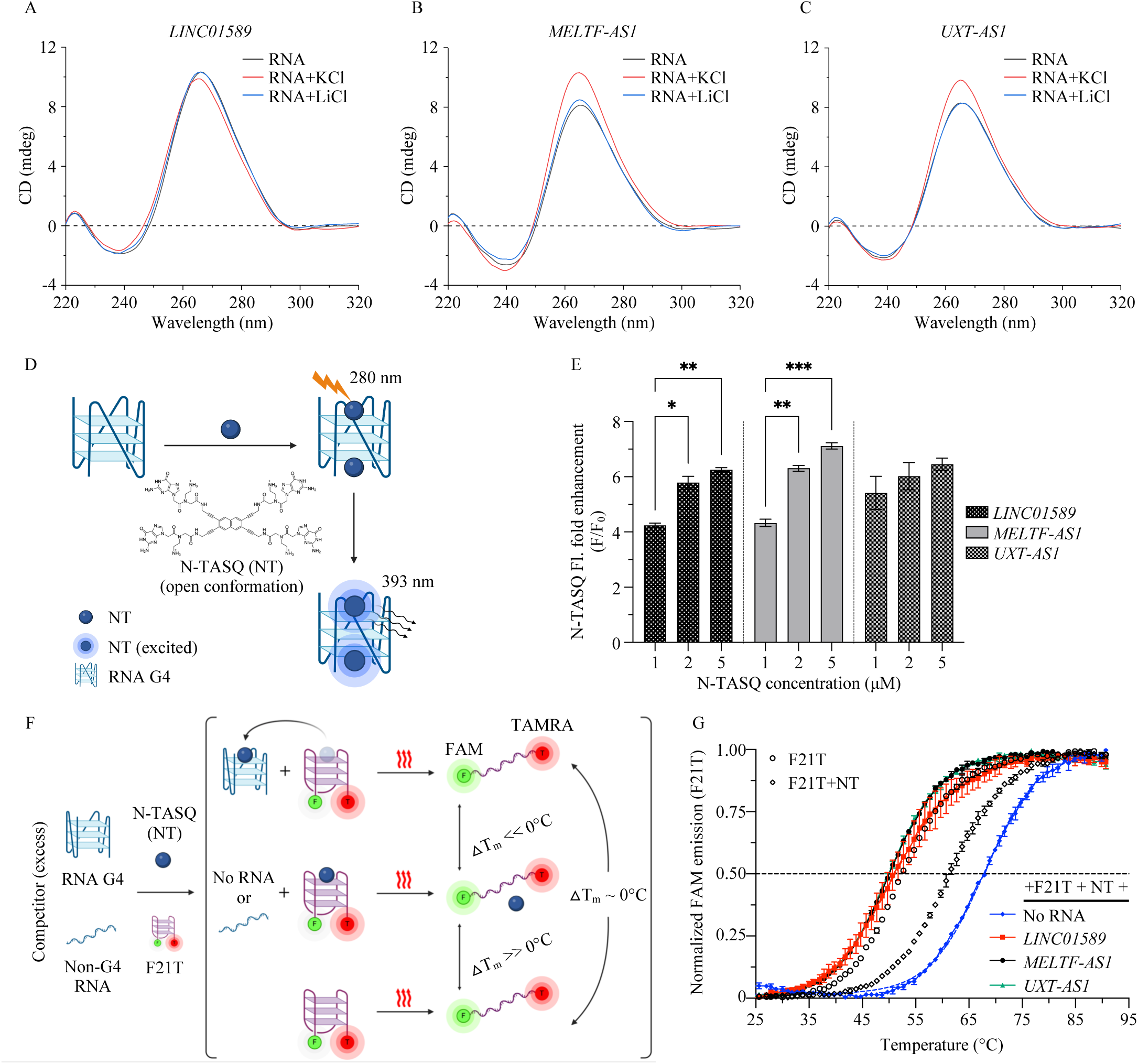
*In vitro* formation of parallel G4s in *LINC01589*, *MELTF-AS1*, and *UXT-AS1* lncRNAs. A-C) CD spectroscopy of IVT RNAs (5 µM) folded in the presence or absence of 100 mM KCl or LiCl. Maxima and minima in mean CD spectra at ca. 265 and 240 nm, respectively, correspond to parallel G4 topologies. D) NaphthoTASQ (N-TASQ or NT) fluorescence enhancement assay of folded synthetic RNAs (2 µM) titrated with N-TASQ (1 – 5 µM). E) Increased fold enhancements in mean ± SD N-TASQ fluorescence (F/ F_0_: N-TASQ fluorescence in the presence of RNA/ absence of RNA) at 393 nm when excited with 280 nm show G4-formation. F) Fluorescence Resonance Energy Transfer-Melting Competition (FRET-MC) assay with folded FAM-21-TAMRA: F21T (0.2 µM) in the presence of N-TASQ (1 µM) and excess of folded synthetic RNAs (3 µM). G) Mean ± SEM of normalized FAM emission at maxima from N-TASQ-stabilized F21T in the presence of RNAs, with increasing temperature, correspond to G4-formation. Dashed lines represent the plots fitted using the Boltzmann sigmoidal model of non-linear regression. *P*-values: *P* ≤ 0.05, *P* ≤ 0.01, *P* ≤ 0.001, and *P* ≤ 0.0001 are denoted with one asterisk (*), two asterisks (**), three asterisks (***), and four asterisks (****), respectively. Non-significant *P*-values are not represented.

A G4-specific fluorescence turn-on ligand, NaphthoTASQ (N-TASQ), was used to confirm the G4-folding *via* fluorescence spectroscopy (Figure 4D).^53^ Upon titrating the RNAs with increasing concentrations of N-TASQ, an increase in fluorescence was observed, which validated the *in vitro* G4-formation in *LINC01589*, *MELTF-AS1*, and *UXT-AS1* lncRNAs (Figure 4E; S1B-D). To further substantiate this, a competitive Fluorescence Resonance Energy Transfer-Melting Competitive (FRET-MC) assay was performed. This technique relies on a competition between an excess of putative G4 and a fluorescently-labelled DNA G4 (F21T) for binding to a G4-ligand (N-TASQ) (Figure 4F).^53–55^ The melting temperature (T_m_) of N-TASQ-stabilized F21T (T_m_ = 67.9 °C) decreased in the presence of *LINC01589*, *MELTF-AS1*, and *UXT-AS1* lncRNAs (ΔT_m_ = −17.3, −18.1, and −18.2 °C, respectively), which supports the formation of G4s in these lncRNAs (Figure 4G).

Furthermore, we performed CD-melting experiments to assess the thermostability of G4s in the supplementation of 10 mM KCl. The obtained T_m_ values (44.7, 56.8, and 65.4 °C for the G4s formed by *LINC01589*, *MELTF-AS1*, and *UXT-AS1* lncRNAs, respectively) are in agreement with stable to highly stable G4s (Figure 5A). The CD-melting spectra of *UXT-AS1* exhibit a biphasic pattern with a modestly weaker first transition, suggesting the presence of two distinct species. Nevertheless, the pattern of melting temperatures for *LINC01589*, *MELTF-AS1*, and *UXT-AS1* lncRNAs is in accordance with the greater thermal stability likely associated with a higher number of G-quartets (2G, 3G, and 4G, respectively).^64,65^

**Figure 5.**
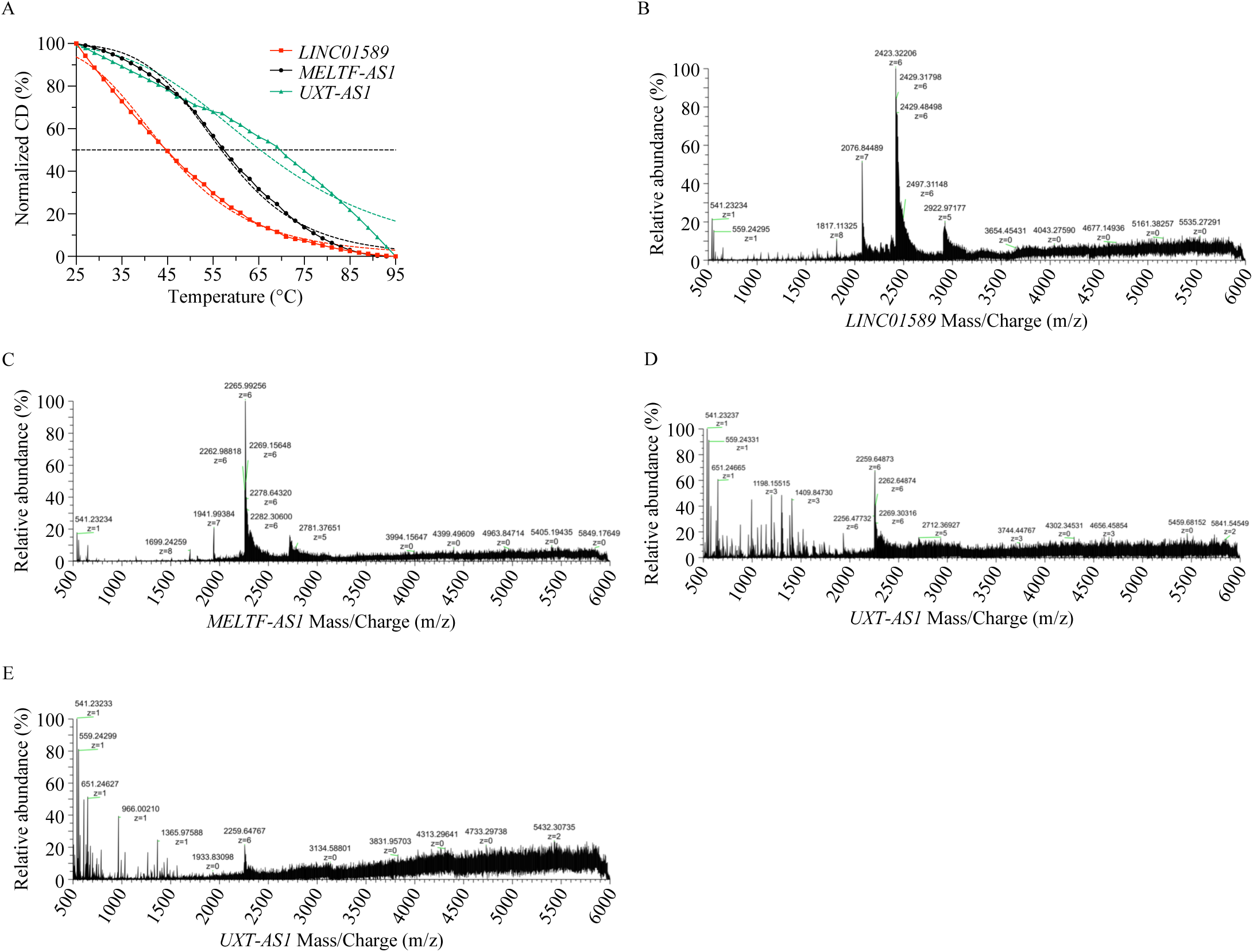
*In vitro* formation of stable and intramolecular G4s in *LINC01589*, *MELTF-AS1*, and *UXT-AS1* lncRNAs. A) Normalized mean CD spectra of folded synthetic RNAs (1 µM) at maxima obtained in the CD spectra, with increasing temperature. Greater thermal stability of RNAs shows an association with a higher number of G-quartets. Dashed lines represent the plots fitted using the dose response-inhibition model of non-linear regression. B-D) Electrospray Ionization Mass Spectrometry (ESI-MS) mass/charge (m/z) spectra of folded synthetic RNAs (10 µM) show highest abundance MS peaks corresponding to the mass of unimolecular or intramolecular G4s. E) ESI-MS m/z spectra of an additional peak of folded synthetic *UXT-AS1* lncRNA (10 µM) with high abundance and significance absorbance at 260 nm in the HILIC chromatogram and UV-Vis data, respectively, shows an MS peak corresponding to the mass of unimolecular or intramolecular G4.

We also performed mass spectrometry analyses with *LINC01589*, *MELTF-AS1*, and *UXT-AS1* lncRNAs folded in 100 mM ammonium acetate solution (pH 7.0) to investigate the molecularity of their G4s.^58,59^ A combination of Hydrophilic Interaction Liquid Chromatography (HILIC) and Electrospray Ionization Mass Spectrometry (ESI-MS) analyses showed that across all the lncRNA G4s tested, the highest abundance MS peaks corresponded to the mass of unimolecular or intramolecular G4s (Figure 5B-D; S2E-J). Notably, for *UXT-AS1*, the ESI-MS analysis of an additional peak with high abundance and significant absorption at 260 nm in the HILIC chromatogram and UV-Vis data, respectively, revealed an MS peak corresponding to the mass of unimolecular or intramolecular G4s (Figure 5E; S2G). This observation is consistent with the CD-melting data for *UXT-AS1*, indicating the presence of two distinct species (Figure 5A). Taken together, this series of *in vitro* results demonstrate the formation of stable, parallel, and intramolecular G4s in *LINC01589*, *MELTF-AS1*, and *UXT-AS1* lncRNAs.

Monovalent cations play a crucial role in the stability and topology of G4s. K^+^ and Na^+^ are commonly employed to enhance G4-stability, while Li^+^ is often used to decrease G4-stability.^61,63^ A 90-, 158-, and 263-fold of Thioflavin T (ThT) fluorescence enhancement was obtained for the *LINC01589*, *MELTF-AS1*, and *UXT-AS1* lncRNAs, respectively, without supplementation of KCl or LiCl (Figure S3A-G).^66^ The supplementation of KCl or LiCl failed to elicit a logarithmic fold-change in G4-stability of the three lncRNAs (Figure S3B-G). Interestingly, the Reverse Transcriptase stop (RT stop) assay indicated that the reverse transcriptions of all lncRNA G4s into full-length cDNAs were modulated by K^+^ and Li^+^, albeit in somewhat erratic manner (Figure S4A-G; Table S3).^67–71^ Collectively, these findings suggest a modest concentration-dependent modulation of G4s formed by these lncRNAs, which warrants a separate investigation.

We next investigated the relative importance of individual G-tracts in each lncRNA towards maintaining the structural integrity of the cognate G4s. The IVT-derived wild-type (WT) lncRNAs and deletion mutants (ι1) of lncRNAs devoid of singular G-tracts were analyzed on 15% native PAGE stained with ThT (Figure S5A-C; Table S3).^72^ For *LINC01589* lncRNA, a faint ThT-stained band was observed only in the ι11 lncRNA when compared to the wild-type lncRNA (Figure S5A, D). In contrast, the band intensities decreased significantly for all deletion mutants of *MELTF-AS1* and *UXT-AS1* lncRNAs, except cognate Δ1, when compared to the wild-type lncRNA (Figure S5B, C, E, F). These results illustrate the importance of the specific G-tract(s) of *LINC01589* (first G-tract), *MELTF-AS1* (second, third, fourth, fifth, and sixth G-tracts), and *UXT-AS1* (second, third, fourth, and fifth G-tracts) towards maintaining the cognate G4-stability.

We further explored the formation of G4s within lncRNAs by a novel approach we have termed Competitor Oligonucleotide Mediated-G4 Disruption (COM-G4D). It involves the incubation of lncRNA G4s with increasing concentrations of competitor DNA oligonucleotides complementary to the full-length G4-forming sequences or different G-tract(s) of lncRNA G4s. The samples were run on 15% native PAGE stained with ThT to visualize the influence of DNA competitors on G4-formation (Figure S9A; Table S3). The increasing concentrations of full-length competitor DNAs for all three lncRNAs (*LINC01589*_Comp FL, *MELTF-AS1*_Comp FL, and *UXT-AS1*_Comp FL) resulted in a significant decrease in the band intensities of respective lncRNA G4s when compared to the no-competitor bands (Figure S9B-D). Further, increasing concentrations of *LINC01589_*Comp 1, *MELTF-AS1*_Comp 2, *UXT-AS1*_Comp 12, and *UXT-AS1*_Comp 5 resulted in a decrease in band intensities (Figure S10A, B, S11A, C, S12A, B, E; Table S3). These competitor DNAs are able to bind their target sites on the lncRNAs and disrupt G4-formation in each case. However, the other competitor DNAs failed to affect similar changes in the band intensities (Figure S10-12). The results observed upon the use of *LINC01589_*Comp 1, *MELTF-AS1*_Comp 2, *UXT-AS1*_Comp 12, and *UXT-AS1*_Comp 5 are in agreement with the results observed for the corresponding deletion mutants. Together, these findings highlight the key G-tracts within these lncRNAs that play a significant role in stabilizing their cognate G4s.

### *LINC01589*, *MELTF-AS1*, and *UXT-AS1* lncRNAs harbour G4s in CRC

We next sought to investigate the *in cella* formation of *LINC01589*, *MELFT-AS1*, and *UXT-AS1* lncRNA G4s in CRC using the HT-29 cell line, in which the expression levels of these lncRNAs are nearly identical, allowing for a standardized analysis (Figure 3A, B). We first performed RNA G4 Immunofluorescence and Fluorescence *in situ* hybridization (RNA G4-Immuno-FISH) by probing the nuclease-treated or untreated HT-29 cells with the G4-specific antibody BG4.^38,39^ This was followed by RNA FISH, where G4-labelled and nuclease-treated or untreated HT-29 cells were hybridized with the DNA probes designed against the lncRNAs.^40–42^ Confocal images were quantified as indicated in the materials and methods section. *NRAS* mRNA was used as a positive control for the RNA G4-Immuno-FISH since it is well-known to harbor G4 *in cella* (Figure 6A; Table S4).^73^

**Figure 6.**
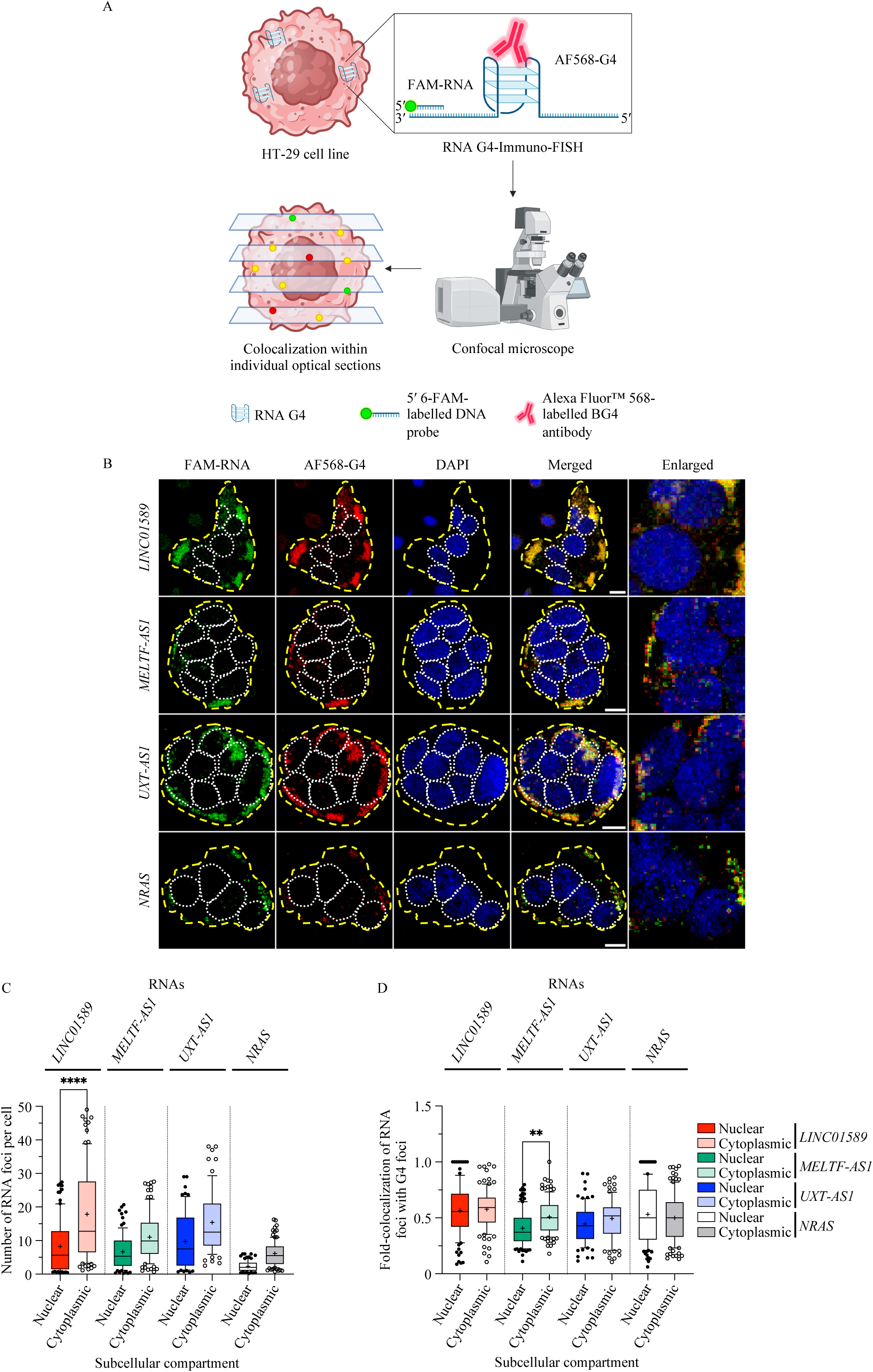
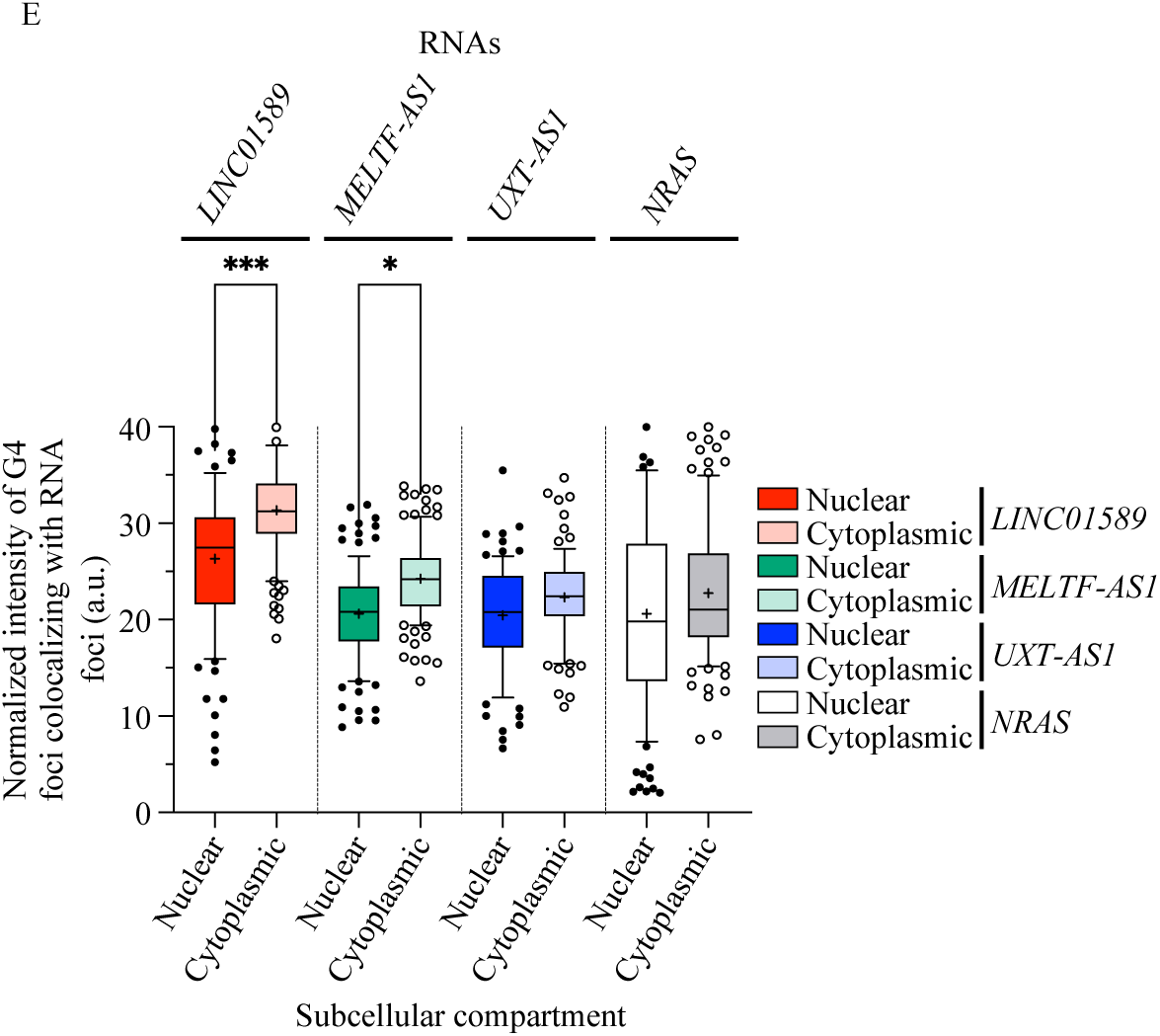
RNA G4 Immunofluorescence and Fluorescence in situ hybridization (RNA G4-Immuno-FISH) identifies the G4s in *LINC01589*, *MELTF-AS1*, and *UXT-AS1* lncRNAs in CRC. A) RNA G4-Immuno-FISH in HT-29 cells probed with G4-specific FLAG-BG4 – anti-FLAG – Alexa Fluor™ 568 (AF568) antibodies, and hybridized with 5ʹ 6-FAM-labelled DNA probes complementary to the flanking regions of the *in vitro* validated G4-forming regions of RNAs. B) Cell clusters (marked with dashed yellow lines) imaged using Confocal microscopy (scale bar: 10 µm) in the FAM-RNA and AF568-G4 panels show the presence of RNAs and G4s, respectively. Merged and enlarged panels show foci corresponding to RNAs and G4s, and colocalized foci corresponding to RNA G4s. Individual optical sections with respect to the nucleus (DAPI staining, marked with dotted white lines) analyzed for C) mean number of RNA foci per cell, D) mean fold-colocalization of the RNA foci with the G4 foci, and E) mean of normalized intensities of G4 foci colocalizing with the RNA foci, in nuclear or cytoplasmic compartment, with whiskers at 10-90 percentile, show the presence of RNA G4s. Figure panels C and D share the legends. *P*-values: *P* ≤ 0.05, *P* ≤ 0.01, *P* ≤ 0.001, and *P* ≤ 0.0001 are denoted with one asterisk (*), two asterisks (**), three asterisks (***), and four asterisks (****), respectively. Non-significant *P*-values are not represented.

The G4 *foci* were observed in the cytoplasmic and nuclear regions of the HT-29 cells, suggesting the presence of DNA and RNA G4s (Figure 6B). *LINC01589 foci* were significantly higher in the cytoplasmic region of HT-29 cells. Furthermore, both *MELTF-AS1* and *UXT-AS1 foci* were also enriched in the cytoplasm over the nucleus (*p* = 0.19 and 0.09, respectively; Figure 6B, C). This suggests the localization of these lncRNAs primarily in the cytoplasm. The positive control *NRAS* mRNA showed higher cytoplasmic localization as expected (*p* = 0.34; Figure 6B, C). Colocalizations of 0.49-to 0.58-fold and 0.41-to 0.57-fold were observed for the lncRNA *foci* with the G4 *foci* in the cytoplasm and nucleus, respectively. However, the observed colocalization for *MELTF-AS1* was significantly higher in the cytoplasm (Figure 6B, D). This finding indicates the presence of G4s in these lncRNAs, but their transient formation may explain the moderate colocalization observed. The increased intensity of the G4 *foci* colocalizing with the *LINC01589* and *MELTF-AS1 foci* in the cytoplasm suggests the increased levels of G4s in the cytoplasmic counterparts of these lncRNAs (Figure 6B, E). Analogous results were observed for the positive control *NRAS* mRNA (Figure 6B, D, E).

Upon DNase or RNase treatment, G4 *foci* were reduced in the nucleus or cytoplasm, respectively, suggesting the formation of both DNA and RNA G4s. While the localization of lncRNA *foci* were unaffected in the presence of DNase, a drastic reduction was observed in the presence of RNase, confirming the binding of DNA probes only to the RNAs. The colocalization between RNA G4s and lncRNAs was unaffected by DNase treatment but became negligible upon RNase treatment, confirming the presence of RNA G4s in these lncRNAs (Figure S13A, B).

For further investigation, we performed G4-RNA-specific Precipitation coupled with the Reverse Transcription-quantitative Polymerase Chain Reaction (G4RP-RT-qPCR) from the HT-29 cell line. The cell lysate was incubated with a biotinylated TASQ (for template-assembled synthetic G-quartets) named BioCyTASQ, which serves as a molecular bait for G4s (Figure 7A).^43^ As a negative control, the cell lysate was incubated with biotin to determine the non-specific interaction of nucleic acids with the biotin moiety or the streptavidin beads. BioCyTASQ was shown to stabilize the *LINC01589*, *MELTF-AS1*, and *UXT-AS1* G4s *in vitro* (Figure S14A-F). Hence, live-cell fixation with paraformaldehyde was performed prior to the cell lysis and incubation of the lysate with BioCyTASQ. The RT-qPCR analysis of the precipitated G4-forming RNAs was carried out using suitable primer pairs (Table S2).^18,43–45^ The fold-change in the level of lncRNAs in BioCyTASQ precipitated samples relative to the input (5%) cell lysate would provide evidence for G4-formation in these lncRNAs. A 2.56-, 0.75-, and 3.30-fold change was observed for the *LINC01589*, *MELFT-AS1*, and *UXT-AS1*, respectively, validating G4-formation in these lncRNAs (Figure 7B). The decreased fold change observed for *MELTF-AS1* could be attributed to the transient formation of G4s.^18^ The positive control *NRAS* mRNA showed a 3.56-fold change, which is in line with reported values (Figure 7B; Table S2).^18,45^ No fold-change (no Ct value) was observed for these lncRNAs and mRNA in biotin samples (negative control), which confirms the absence of non-specific interactions (Figure 7B). Together, these results successfully establish the presence of G4s in the *LINC01589*, *MELFT-AS1*, and *UXT-AS1* lncRNAs in the CRC.

**Figure 7.**
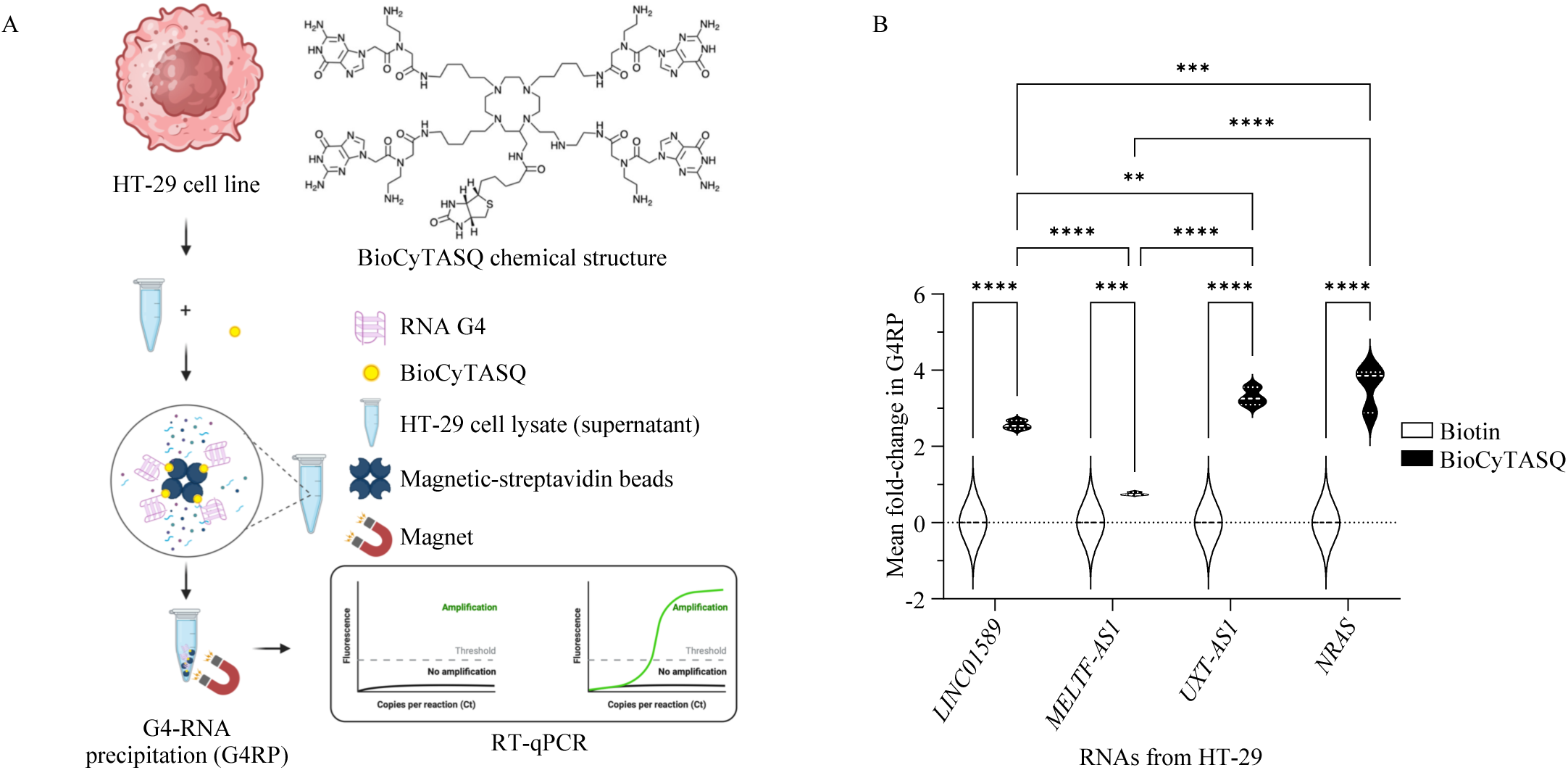
G4-RNA-specific precipitation coupled with the Reverse Transcription-quantitative Polymerase Chain Reaction (G4RP-RT-qPCR) validates the G4s in *LINC01589*, *MELTF-AS1*, and *UXT-AS1* lncRNAs in CRC. A) G4RP-RT-qPCR from HT-29 cells, including RNA precipitation from cell lysate incubated with BioCyTASQ, using magnetic-streptavidin beads, and RT-qPCR of precipitated RNAs in the presence of suitable primer pairs. B) Mean ± SEM fold-change in the levels of RNAs [5 * {2^(Mean Ct input – Ct G4RP or biotin)^}] indicate the presence of RNA G4s. *P*-values: *P* ≤ 0.05, *P* ≤ 0.01, *P* ≤ 0.001, and *P* ≤ 0.0001 are denoted with one asterisk (*), two asterisks (**), three asterisks (***), and four asterisks (****), respectively. Non-significant *P*-values are not represented.

## Conclusion

Advancements in high-throughput techniques for analyzing human transcriptome have revealed the involvement of lncRNAs in key biological processes and their association with various pathophysiological conditions.^74^ Despite the established connection between lncRNA dysregulation and diseases, such as cancers, the role of their secondary structures in modulating disease progression remains relatively unexplored.^7–10^ Although CRC ranks high in cancer-related mortalities, not much is known about the molecular basis of its carcinogenesis.^23^ Successful treatment of CRC by chemo-and immuno-therapy as well as surgery resides in early-stage detection.^24^ This highlights the pressing need for identifying biomarkers capable of early diagnosis. The tissue-specific expression and presence in various bodily fluids make lncRNAs promising candidates.^29,30^ This explains why, over the last ten years, numerous studies have focused on dysregulated lncRNAs in CRC.^26,27,31^

LncRNAs harbor G4-prone sequences, and these G4s are likely to play a significant role in lncRNA biology and disease progression.^19^ Recently, a few G4-harboring lncRNAs have been linked with CRC modulation, highlighting the possible roles of G4s as hubs for various CRC-associated regulatory entities.^32–34^ However, the lack of understanding regarding G4-formation in other dysregulated lncRNAs and the overall influence of their G4s on CRC progression led us to explore the G4-formation in additional CRC-dysregulated lncRNAs. Using *in silico* tools and *in vitro* biophysical techniques, we demonstrate that CRC-dysregulated *LINC01589*, *MELFT-AS1*, and *UXT-AS1* lncRNAs can form stable, parallel, and intramolecular G4s. While the observed pattern of thermostability and *in vitro* fluorescence enhancements suggests the likely constitution of *LINC01589*, *MELTF-AS1*, and *UXT-AS1* G4s as 2G, 3G, and 4G-G4, respectively, which highlights their structural complexity, further structural investigations are necessary to validate these findings.^64,65^ We also highlight the expression of these lncRNAs in cancers other than CRC.

Our results show that the G4s formed by *LINC01589*, *MELTF-AS1*, and *UXT-AS1* lncRNAs are present *in cella* in the context of CRC. All three lncRNAs are prevalent in the cytoplasm, but their presence in the nucleus suggests a multifaceted implication in cellular biology. The moderate detection of these lncRNA G4s in both cytoplasmic and nuclear compartments highlights their transient or dynamic formation, likely finely regulated by cytoplasmic and nuclear helicases and chaperones.^18^

LncRNAs, being rich in secondary structures, possess multiple dynamic conformations.^6,75,76^ Reports suggest an equilibrium between RNA G4s and other secondary structures, including stem-loop and hairpin, within a few RNAs. Such equilibrium is monitored by ion-dependent structural shifts and ligand-dependent G4-modulation, and regulates diverse cellular processes.^77,78^ Collectively, the ability of these lncRNAs to be present across the cell, along with their dynamism, suggests specific functions.

Our results show that the lncRNAs studied here are globally expressed in CRC, and this abundance correlates with that of the corresponding G4s, detected by both qualitative (*i.e.*, optical imaging) and quantitative methods (*i.e.*, G4RP-RT-qPCR). The sole dissonant note is *MELTF-AS1* G4, for which G4RP-RT-qPCR and RNA G4-Immuno-FISH results do not correlate. This might originate in its stability, transient nature, or the fact that specific chaperones and G4-binding proteins may hinder the binding of BioCyTASQ. Further studies will thus be required to understand this discrepancy thoroughly.

Collectively, our results contribute to a better understanding of the functional relevance of lncRNAs in CRC, their ability to bear stable G4-motifs, their atypical cellular localization, along with their expression in cancers other than CRC. They thus point to a possible involvement in the regulation of cellular pathways in different types of cancer. Our results also open the way toward using lncRNA G4s to isolate them and identify their interacting partners, proteins or nucleic acids. Indeed, considering the length of the three lncRNAs, it is evident that the G4s are not the only structural motif involved in interactions with proteins and other nucleic acids. Therefore, using G4s as a convenient handle for isolating hitherto unexplored lncRNAs will undoubtedly advance our understanding of CRC biology and progression.

## Supporting information

Supplementary word and excel

## Data Availability

The data is available from the author upon request. The ImageJ macro for the analyses of confocal microscopy images is available on GitHub (https://github.com/ICMUB/FISH-lncRNA-IF-BG4).

## Acknowledgments

The authors thank the Department of Biological Sciences and Engineering, Department of Chemistry, Common Research & Technology Development Hub, and the Central Instrumentation facility at Indian Institute of Technology Gandhinagar, Gandhinagar, Gujarat, India; the Polyamines & Porphyrins: Development & Applications group, SATT Sayens, and the PACSMUB platform at Institut de Chimie Moléculaire de l’Université de Bourgogne, UMR CNRS 6302, Dijon, France, for providing instrumentation facility. The authors also thank Efftesum Rehman for technical assistance with the RT stop assay. All the illustrations are Created with BioRender.com.

---

**Figure S1. *In vitro* formation of parallel G4s in *LINC01589*, *MELTF-AS1*, and *UXT-AS1* lncRNAs.** A) CD spectroscopy of folded synthetic RNAs (1 µM). Maxima and minima in mean CD spectra at ca. 265 and 240 nm, respectively, correspond to parallel G4 topologies. B-D) N-TASQ fluorescence enhancement assay of synthetic RNAs (2 µM) folded in 10 mM Tris-HCl (pH 7.5) and 0.1 mM EDTA (pH 8.0), and titrated with N-TASQ (1 – 5 µM). Increased mean fluorescence emission spectra when excited at 280 nm show G4-formation.

**Figure S2. *LINC01589*, *MELTF-AS1*, and *UXT-AS1* lncRNAs forms intramolecular G4s.** A-C) N-TASQ fluorescence enhancement assay of synthetic RNAs (2 µM) folded in 100 mM Ammonium acetate solution (pH 7.0), and titrated with N-TASQ (1 – 5 µM). Increased mean fluorescence emission spectra when excited at 280 nm show G4-formation. D) FRET-MC assay with folded F21T (0.2 µM) in the presence of N-TASQ (1 µM) and excess of synthetic RNAs (3 µM) folded in 100 mM ammonium acetate solution (pH 7.0). Mean ± SEM of normalized FAM emission at maxima from N-TASQ-stabilized F21T in the presence of RNAs, with increasing temperature, correspond to G4-formation. E-G) Hydrophilic Interaction Liquid Chromatography (HILIC) chromatogram of folded synthetic RNAs (10 µM) with relative abundance and absorbance at 260 nm. H-J) Deconvoluted mass of RNAs with highest abundance MS peaks corresponding to the mass of unimolecular or intramolecular G4s.

**Figure S3. Specific monovalent cations affect the stability of *LINC01589*, *MELTF-AS1*, and *UXT-AS1* lncRNA G4s.** A) Thioflavin T (ThT) fluorescence enhancement assay of IVT RNAs (2 µM) folded in the presence or absence of 100 mM KCl or LiCl with ThT (2 µM). B-D) Increased mean fluorescence emission spectra when excited at 445 nm, and E-G) increased fold enhancements in mean ± SEM ThT fluorescence (F/ F0: ThT fluorescence in the presence of RNA/ absence of RNA) at 488 nm when excited with 445 nm correspond to G4-formation in the presence or absence of specific monovalent cations (K^+^ or Li^+^). *P*-values: *P* ≤ 0.05, *P* ≤ 0.01, *P* ≤ 0.001, and *P* ≤ 0.0001 are denoted with one asterisk (*), two asterisks (**), three asterisks (***), and four asterisks (****), respectively. Non-significant *P*-values are not represented.

**Figure S4. Reverse Transcriptase stop (RT stop) assay validates the effect of specific monovalent cations on the stability of *LINC01589*, *MELTF-AS1*, and *UXT-AS1* lncRNA G4s.** A) RT stop assay of IVT RNAs (2 µM) folded in the presence of 0 – 150 mM KCl or LiCl, using 5ʹ Texas Red (Tx Red)-labelled primer (100 nM) binding to the primer binding site in IVT RNAs, and M-MLV RT. The presence of G4 in the RNA strand stalls RT, precluding the transcription of RNA into full-length cDNA, resulting in a very weak or negligible intensity full-length cDNA band in denaturing PAGE (15%). B-G) Denaturing PAGE show bands (*) corresponding to full-length cDNA in RT stop assay of RNAs in the presence of B-D) KCl and E-G) LiCl. Mean ± SD of their normalized intensities indicates the stabilization or destabilization of G4s in the presence of specific monovalent cations (K^+^ or Li^+^). ssDNA ladder indicates the corresponding band size. *P*-values: *P* ≤ 0.05, *P* ≤ 0.01, *P* ≤ 0.001, and *P* ≤ 0.0001 are denoted with one asterisk (*), two asterisks (**), three asterisks (***), and four asterisks (****), respectively. Non-significant *P*-values are not represented.

**Figure S5. G-tracts influence the stability of *LINC01589*, *MELTF-AS1*, and *UXT-AS1* lncRNA G4s.** A-C) Wild-type (WT) RNAs and deletion mutants (Δ) of RNAs devoid of singular G-tracts synthesized using IVT. D-F) Native PAGE (15%) of folded IVT-derived wild-type and deletion mutants of RNAs (2 µM) stained with ThT (0.5 µM) show bands (*) corresponding to cognate G4s. Mean ± SD of their normalized intensities in each mutant indicates G4-formation. *P*-values: *P* ≤ 0.05, *P* ≤ 0.01, *P* ≤ 0.001, and *P* ≤ 0.0001 are denoted with one asterisk (*), two asterisks (**), three asterisks (***), and four asterisks (****), respectively. Non-significant *P*-values are not represented.

**Figure S6. First G-tract is crucial for the stability of *LINC01589* lncRNA G4.** A-F) ThT fluorescence enhancement assay of wild-type (WT) and deletion mutants (Δ) of RNAs (2 µM) folded in the presence or absence of 100 mM KCl or LiCl with ThT (2 µM). A-C) Increased mean fluorescence emission spectra, and D-F) fold enhancements in mean ± SEM ThT fluorescence at 488 nm when excited with 445 nm correspond to G4-formation in the presence or absence of specific monovalent cations (K^+^ or Li^+^). *P*-values: 0.0332 (*), 0.0021 (**), 0.0002 (***), <0.0001 (****). Non-significant P-values are not represented.

**Figure S7. Second, third, fourth, fifth, and sixth G-tracts are crucial for the stability of *MELTF-AS1* lncRNA G4.** A-F) ThT fluorescence enhancement assay of wild-type (WT) and deletion mutants (Δ) of RNAs (2 µM) folded in the presence or absence of 100 mM KCl or LiCl with ThT (2 µM). A-C) Increased mean fluorescence emission spectra, and D-F) fold enhancements in mean ± SEM ThT fluorescence at 488 nm when excited with 445 nm correspond to G4-formation in the presence or absence of specific monovalent cations (K^+^ or Li^+^). *P*-values: 0.0332 (*), 0.0021 (**), 0.0002 (***), <0.0001 (****). Non-significant P-values are not represented.

**Figure S8. Second, third, fourth, and fifth G-tracts are crucial for the stability of *UXT-AS1* lncRNA G4.** A-F) ThT fluorescence enhancement assay of wild-type (WT) and deletion mutants (Δ) of RNAs (2 µM) folded in the presence or absence of 100 mM KCl or LiCl with ThT (2 µM). A-C) Increased mean fluorescence emission spectra, and D-F) fold enhancements in mean ± SEM ThT fluorescence at 488 nm when excited with 445 nm correspond to G4-formation in the presence or absence of specific monovalent cations (K^+^ or Li^+^). *P*-values: 0.0332 (*), 0.0021 (**), 0.0002 (***), <0.0001 (****). Non-significant P-values are not represented.

**Figure S9. Competitor Oligonucleotide Mediated-G4 Disruption (COM-G4D) indicating the loss of *LINC01589*, *MELTF-AS1*, and *UXT-AS1* lncRNA G4s.** A) COM-G4D using the competitor DNA oligonucleotides complementary to full-length G4-forming sequences of RNAs, binding to the cognate sequences, and blocking the participation of all G-tracts in G4-formation. B-D) Native PAGE (15%) of IVT RNAs (2 µM) folded in the presence of competitor DNA oligonucleotides (0 - 2.4 mol. eq.) and stained with ThT (0.5 µM) show bands (*) corresponding to G4s. Mean ± SD of their normalized intensities in the presence of competitors indicates G4-formation. *P*-values: *P* ≤ 0.05, *P* ≤ 0.01, *P* ≤ 0.001, and *P* ≤ 0.0001 are denoted with one asterisk (*), two asterisks (**), three asterisks (***), and four asterisks (****), respectively. Non-significant *P*-values are not represented.

**Figure S10. COM-G4D on the G-tracts of *LINC01589* lncRNA G4.** A) COM-G4D using the competitor DNA oligonucleotides complementary to different G-tracts of RNA, binding to the cognate G-tract(s), and blocking their participation in G4-formation. B-E) Native PAGE (15%) of IVT RNAs (2 µM) folded in the presence of competitor DNA oligonucleotides (0 - 2.4 mol. eq.) and stained with ThT (0.5 µM) show bands (*) corresponding to G4s. Mean ± SD of their normalized intensities in the presence of each competitor indicates G4-formation. *P*-values: *P* ≤ 0.05, *P* ≤ 0.01, *P* ≤ 0.001, and *P* ≤ 0.0001 are denoted with one asterisk (*), two asterisks (**), three asterisks (***), and four asterisks (****), respectively. Non-significant *P*-values are not represented.

**Figure S11. COM-G4D on the G-tracts of *MELTF-AS1* lncRNA G4.** A) COM-G4D using the competitor DNA oligonucleotides complementary to different G-tracts of RNA, binding to the cognate G-tract(s), and blocking their participation in G4-formation. B-E) Native PAGE (15%) of IVT RNAs (2 µM) folded in the presence of competitor DNA oligonucleotides (0 - 2.4 mol. eq.) and stained with ThT (0.5 µM) show bands (*) corresponding to G4s. Mean ± SD of their normalized intensities in the presence of each competitor indicates G4-formation. *P*-values: *P* ≤ 0.05, *P* ≤ 0.01, *P* ≤ 0.001, and *P* ≤ 0.0001 are denoted with one asterisk (*), two asterisks (**), three asterisks (***), and four asterisks (****), respectively. Non-significant *P*-values are not represented.

**Figure S12. COM-G4D on the G-tracts of *UXT-AS1* lncRNA G4.** A) COM-G4D using the competitor DNA oligonucleotides complementary to different G-tracts of RNA, binding to the cognate G-tract(s), and blocking their participation in G4-formation. B-E) Native PAGE (15%) of IVT RNAs (2 µM) folded in the presence of competitor DNA oligonucleotides (0 - 2.4 mol. eq.) and stained with ThT (0.5 µM) show bands (*) corresponding to G4s. Mean ± SD of their normalized intensities in the presence of each competitor indicates G4-formation. *P*-values: *P* ≤ 0.05, *P* ≤ 0.01, *P* ≤ 0.001, and *P* ≤ 0.0001 are denoted with one asterisk (*), two asterisks (**), three asterisks (***), and four asterisks (****), respectively. Non-significant *P*-values are not represented.

**Figure S13. RNA G4-Immuno-FISH identifies the G4s in *LINC01589*, *MELTF-AS1*, and *UXT-AS1* lncRNAs in CRC.** A-B) RNA G4-Immuno-FISH in HT-29 cells treated with A) DNase, and B) RNase, probed with G4-specific FLAG-BG4 – anti-FLAG – Alexa Fluor 568™ (AF568) antibodies, and hybridized with 5ʹ 6-FAM-labelled DNA probes. Cell clusters (marked with dashed yellow lines) imaged using Confocal microscopy (scale bar: 10 µm) in the FAM-RNA and AF568-G4 panels show the binding of 6-FAM-labelled DNA probes to the RNAs rather than DNAs, and the presence of G4s, respectively. Merged panel shows foci corresponding to RNAs and G4s, and colocalized foci corresponding to RNA G4s. DAPI staining indicates the nucleus (marked with white dotted lines).

**Figure S14. BioCyTASQ used in G4RP-RT-qPCR stabilizes the *LINC01589*, *MELTF-AS1*, and *UXT-AS1* lncRNA G4s *in vitro*.** A-C) CD spectroscopy of folded synthetic RNAs (1 µM) titrated with BioCyTASQ (1 – 5 µM). No change in mean CD spectra corresponds to no effect of BioCyTASQ on the parallel G4 topologies. D-F) Normalized mean CD spectra of folded synthetic RNAs (1 µM) in the presence of BioCyTASQ (5 µM) at maxima obtained in the CD spectra, with increasing temperature. Greater thermal stability of RNAs in the presence of BioCyTASQ shows its stabilizing effect on the G4s. Dashed lines represent the plots fitted using the dose response-inhibition model of non-linear regression. *P*-values: *P* ≤ 0.05, *P* ≤ 0.01, *P* ≤ 0.001, and *P* ≤ 0.0001 are denoted with one asterisk (*), two asterisks (**), three asterisks (***), and four asterisks (****), respectively. Non-significant *P*-values are not represented.

**Table S1. Details of CRC-dysregulated lncRNAs and their in silico G4-forming potential.**

**Table S2. Forward and reverse primer sequences used for RT-qPCR.**

**Table S3. Details of the DNA and RNA oligonucleotide sequences used for FRET-MC, RT stop assay primer, templates for the IVT of wild-type (WT) and deletion mutants (**Δ) **of RNA PQS, synthetic wild-type (WT) RNA PQS, and competitor DNA oligonucleotides.** cDNA antisense (AS) of RNA PQS and RT stop primer binding site in IVT template are highlighted in bold and underlined, respectively. PQS/ G4-forming regions in synthetic RNAs are marked in red.

**Table S4. 5ʹ 6-FAM-labelled DNA probe sequences used for RNA G4-Immuno-FISH.**

