## Supplementary word and excel for "G-quadruplex formation in long non-coding RNAs dysregulated in colorectal cancer": SI figures_LncRNA G4-CRC manuscript_Submitted to bioRxiv_05.07.2024_SS.pdf

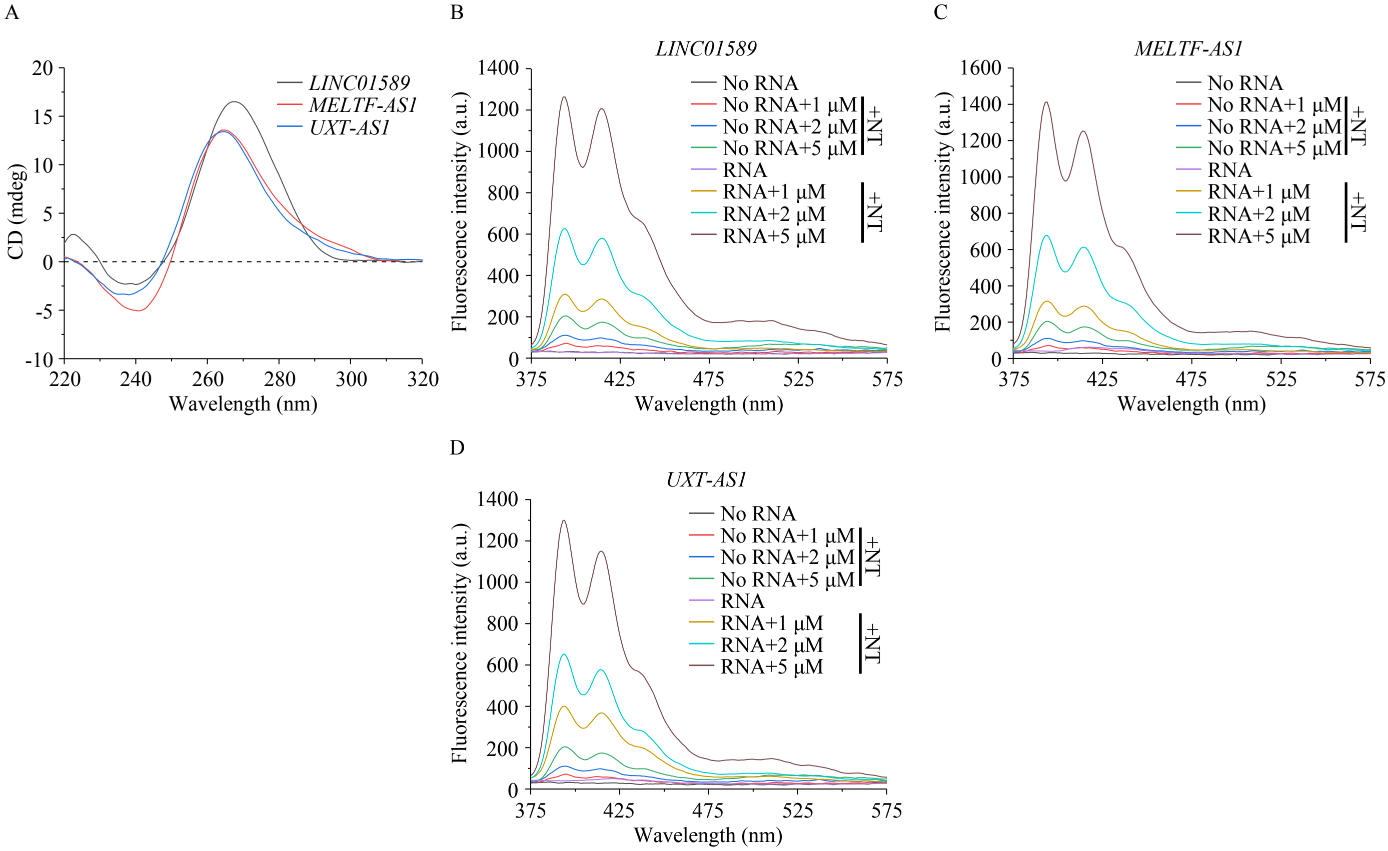


**Figure S1. *In vitro* formation of parallel G4s in *LINC01589*, *MELTF-AS1*, and *UXT-AS1* lncRNAs.** A) CD spectroscopy of folded synthetic RNAs (1 μM). Maxima and minima in mean CD spectra at ca. 265 and 240 nm, respectively, correspond to parallel G4 topologies. B-D) N-TASQ fluorescence enhancement assay of synthetic RNAs (2 μM) folded in 10 mM Tris-HCl (pH 7.5) and 0.1 mM EDTA (pH 8.0), and titrated with N-TASQ (1 – 5 μM). Increased mean fluorescence emission spectra when excited at 280 nm show G4-formation.

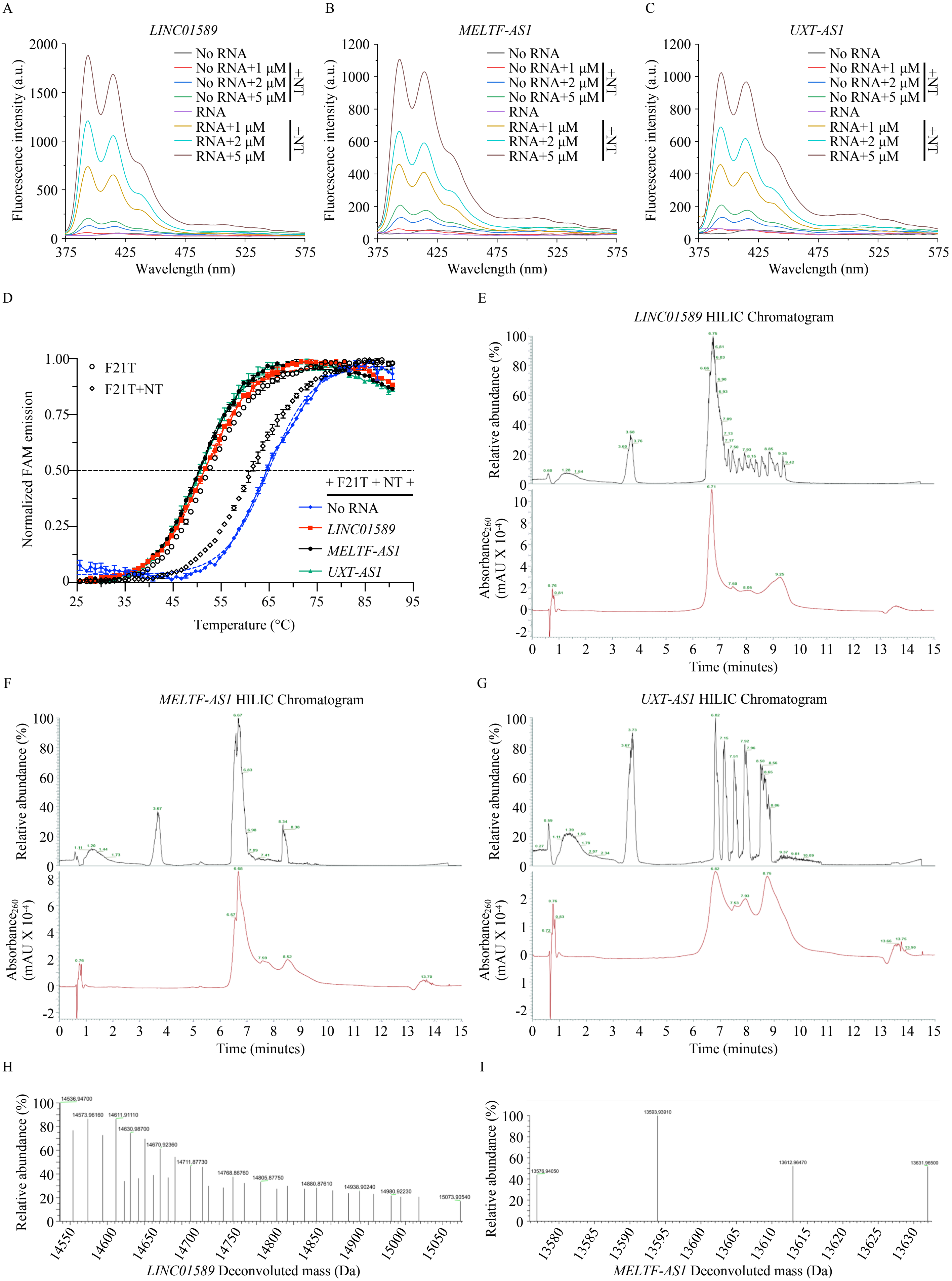
A
LINC01589
Fluorescence intensity (a.u.)
Wavelength (nm)
No RNA
No RNA+1 μM
No RNA+2 μM
No RNA+5 μM
RNA
RNA+1 μM
RNA+2 μM
RNA+5 μM
+NT
B
MELTF-AS1
C
UXT-AS1
D
Normalized FAM emission
Temperature (°C)
F21T
F21T+NT
+ F21T + NT +
No RNA
LINC01589
MELTF-AS1
UXT-AS1
E
LINC01589 HILIC Chromatogram
Relative abundance (%)
Absorbance260 (mAU X 10-4)
Time (minutes)
F
MELTF-AS1 HILIC Chromatogram
G
UXT-AS1 HILIC Chromatogram
H
LINC01589 Deconvoluted mass (Da)
I
MELTF-AS1 Deconvoluted mass (Da)

J

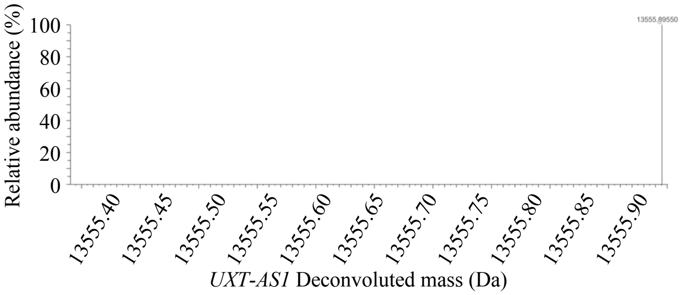


**Figure S2. *LINC01589*, *MELTF-AS1*, and *UXT-AS1* lncRNAs forms intramolecular G4s.** A-C) N-TASQ fluorescence enhancement assay of synthetic RNAs (2 µM) folded in 100 mM Ammonium acetate solution (pH 7.0), and titrated with N-TASQ (1 – 5 µM). Increased mean fluorescence emission spectra when excited at 280 nm show G4-formation. D) FRET-MC assay with folded F21T (0.2 µM) in the presence of N-TASQ (1 µM) and excess of synthetic RNAs (3 µM) folded in 100 mM ammonium acetate solution (pH 7.0). Mean ± SEM of normalized FAM emission at maxima from N-TASQ-stabilized F21T in the presence of RNAs, with increasing temperature, correspond to G4-formation. E-G) Hydrophilic Interaction Liquid Chromatography (HILIC) chromatogram of folded synthetic RNAs (10 µM) with relative abundance and absorbance at 260 nm. H-J) Deconvoluted mass of RNAs with highest abundance MS peaks corresponding to the mass of unimolecular or intramolecular G4s.

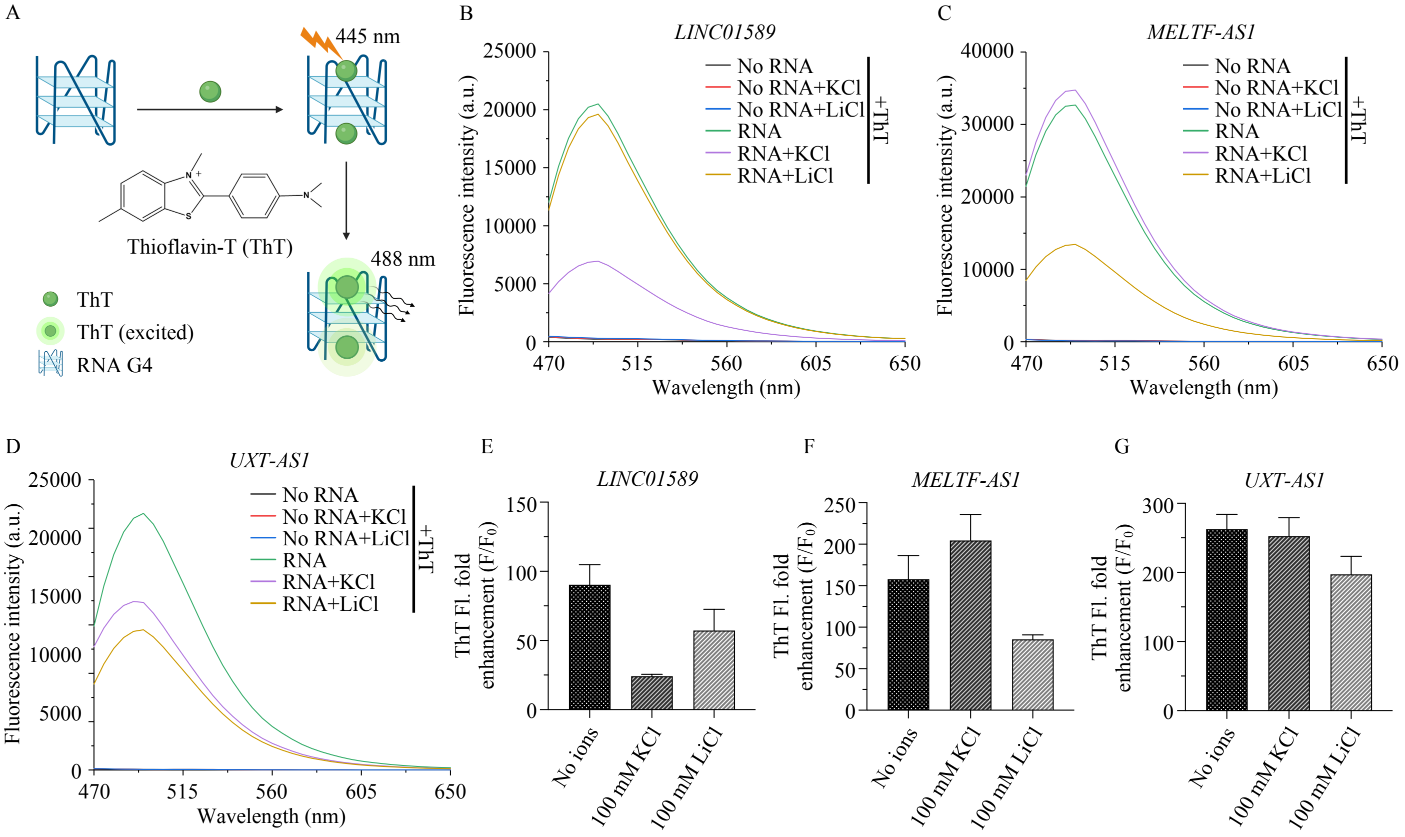


**Figure S3. Specific monovalent cations affect the stability of *LINC01589*, *MELTF-AS1*, and *UXT-AS1* lncRNA G4s.** A) Thioflavin T (ThT) fluorescence enhancement assay of IVT RNAs (2 μM) folded in the presence or absence of 100 mM KCl or LiCl with ThT (2 μM). B-D) Increased mean fluorescence emission spectra when excited at 445 nm, and E-G) increased fold enhancements in mean ± SEM ThT fluorescence (F/ F0: ThT fluorescence in the presence of RNA/ absence of RNA) at 488 nm when excited with 445 nm correspond to G4-formation in the presence or absence of specific monovalent cations ($K^+$ or $Li^+$). *P*-values: $P \leq 0.05$, $P \leq 0.01$, $P \leq 0.001$, and $P \leq 0.0001$ are denoted with one asterisk (*), two asterisks (**), three asterisks (***), and four asterisks (****), respectively. Non-significant *P*-values are not represented.

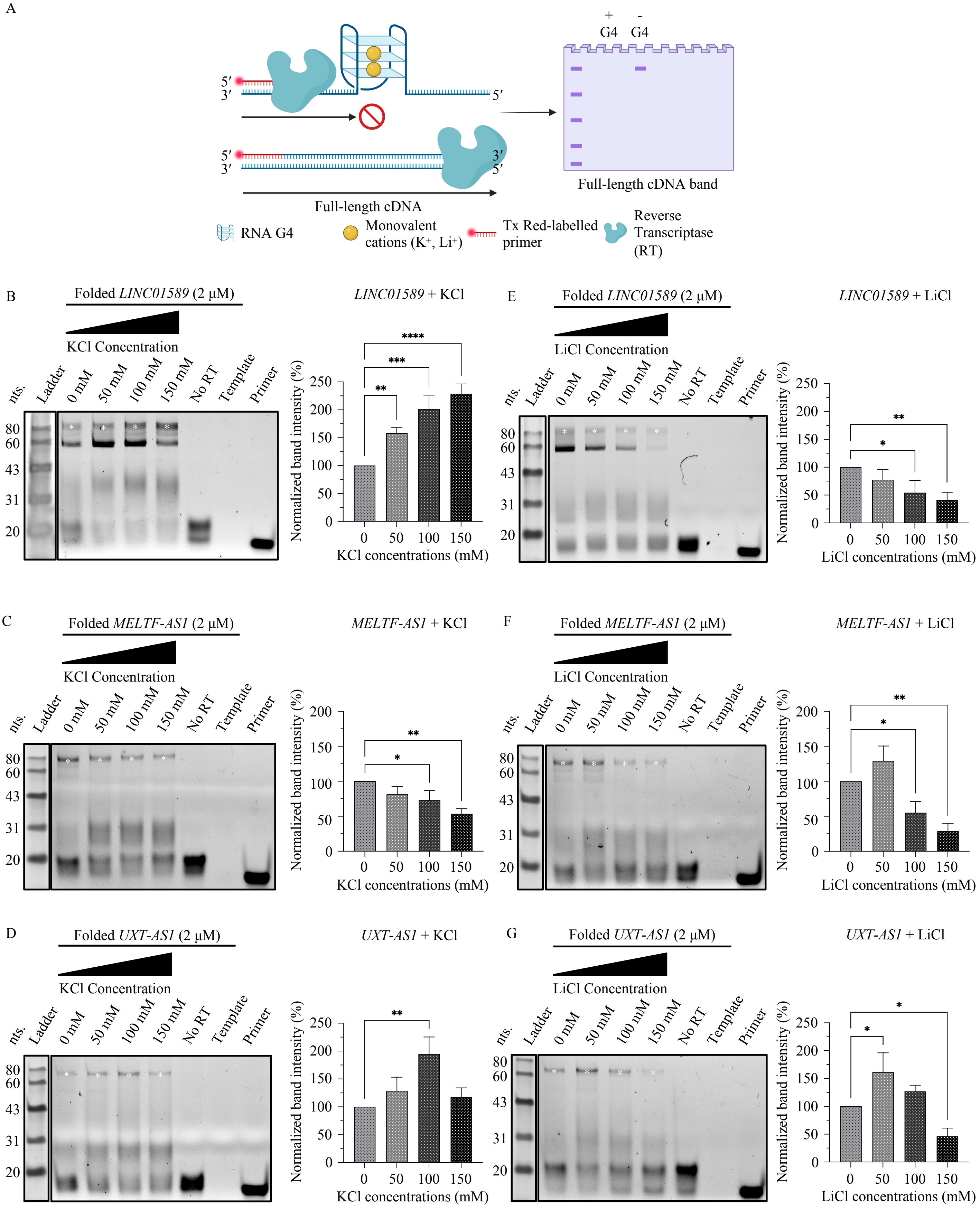


**Figure S4. Reverse Transcriptase stop (RT stop) assay validates the effect of specific monovalent cations on the stability of *LINC01589*, *MELTF-AS1*, and *UXT-AS1* lncRNA G4s.** A) RT stop assay of IVT RNAs (2 µM) folded in the presence of 0 – 150 mM KCl or LiCl, using 5′ Texas Red (Tx Red)-labelled primer (100 nM) binding to the primer binding site in IVT RNAs, and M-MLV RT. The presence of G4 in the RNA strand stalls RT, precluding the transcription of RNA into full-length cDNA, resulting in a very weak or negligible intensity full-length cDNA band in denaturing PAGE (15%). B-G) Denaturing PAGE show bands (*) corresponding to full-length cDNA in RT stop assay of RNAs in the presence of B-D) KCl and E-G) LiCl. Mean ± SD of their normalized intensities indicates the stabilization or destabilization of G4s in the presence of specific monovalent cations ($K^+$ or $Li^+$). ssDNA ladder indicates the corresponding band size. *P*-values: $P \leq 0.05$, $P \leq 0.01$, $P \leq 0.001$, and $P \leq 0.0001$ are denoted with one asterisk (*), two asterisks (**), three asterisks (***), and four asterisks (****), respectively. Non-significant *P*-values are not represented.

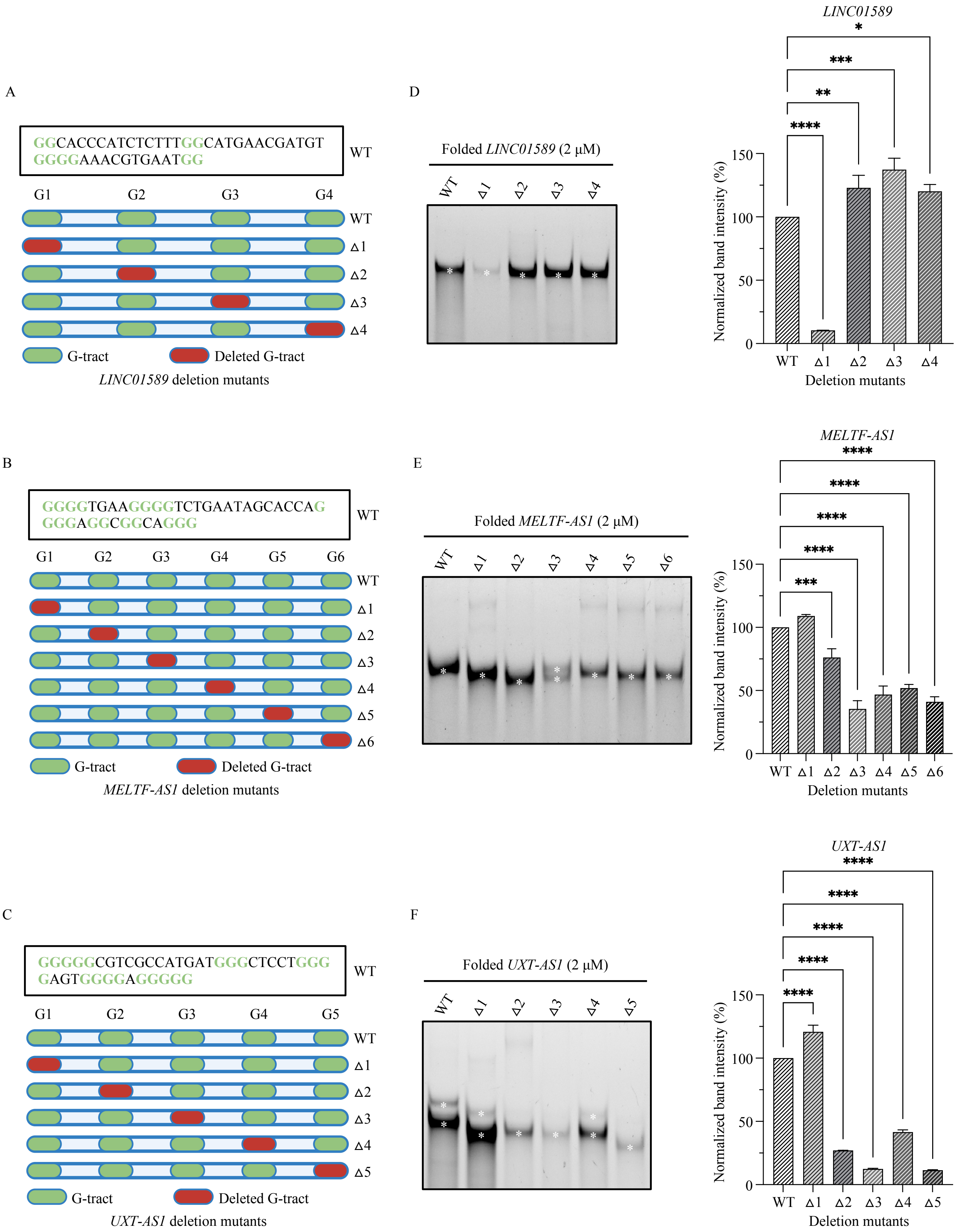


**Figure S5. G-tracts influence the stability of *LINC01589*, *MELTF-AS1*, and *UXT-AS1* lncRNA G4s.** A-C) Wild-type (WT) RNAs and deletion mutants (Δ) of RNAs devoid of singular G-tracts synthesized using IVT. D-F) Native PAGE (15%) of folded IVT-derived wild-type and deletion mutants of RNAs (2 µM) stained with ThT (0.5 µM) show bands (*) corresponding to cognate G4s. Mean ± SD of their normalized intensities in each mutant indicates G4-formation. *P*-values: $P \leq 0.05$, $P \leq 0.01$, $P \leq 0.001$, and $P \leq 0.0001$ are denoted with one asterisk (*), two asterisks (**), three asterisks (***), and four asterisks (****), respectively. Non-significant *P*-values are not represented.

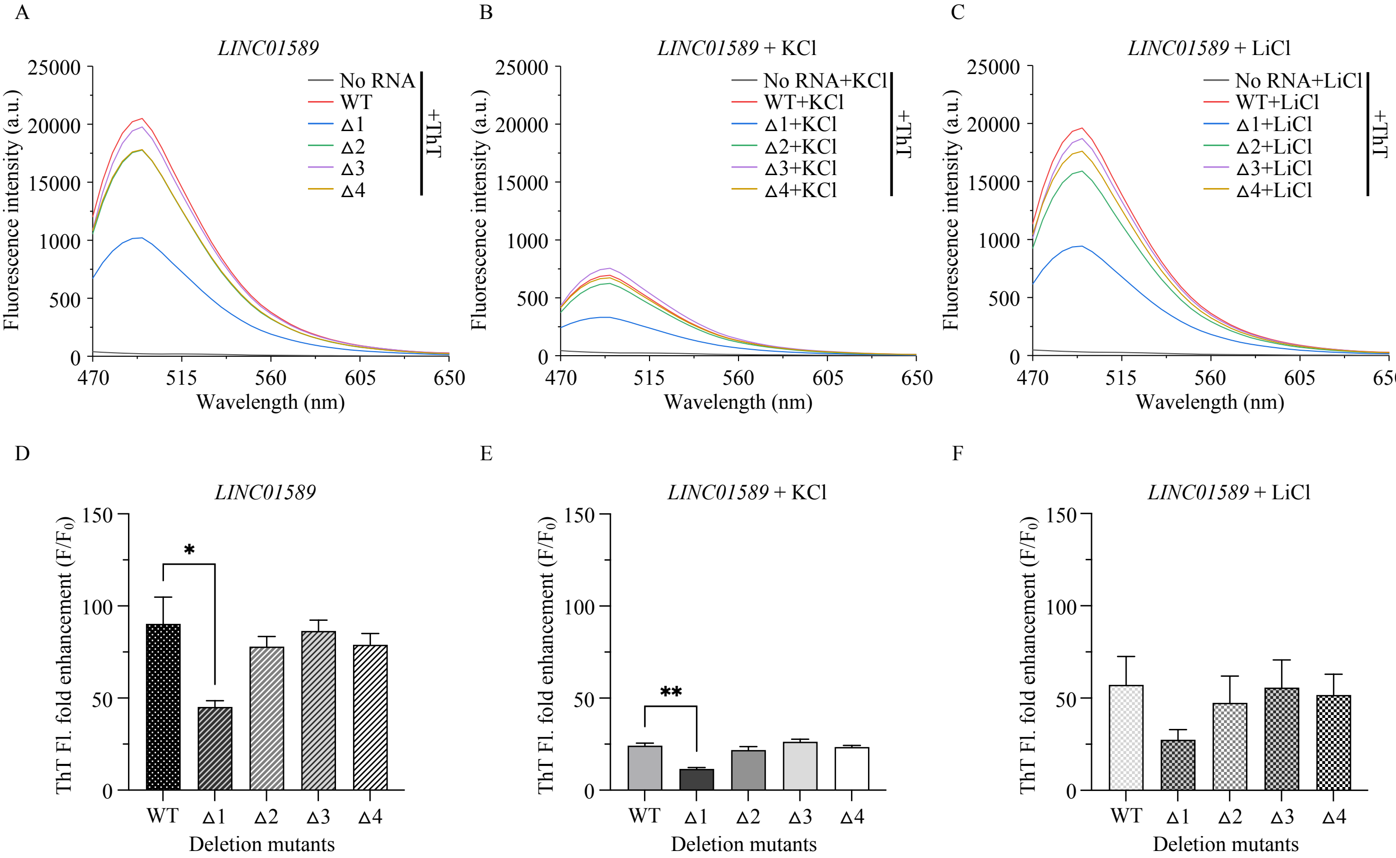


**Figure S6. First G-tract is crucial for the stability of *LINC01589* lncRNA G4.** A-F) ThT fluorescence enhancement assay of wild-type (WT) and deletion mutants (Δ) of RNAs (2 μM) folded in the presence or absence of 100 mM KCl or LiCl with ThT (2 μM). A-C) Increased mean fluorescence emission spectra, and D-F) fold enhancements in mean ± SEM ThT fluorescence at 488 nm when excited with 445 nm correspond to G4-formation in the presence or absence of specific monovalent cations ($K^+$ or $Li^+$). *P*-values: 0.0332 (*), 0.0021 (**), 0.0002 (***), <0.0001 (****). Non-significant P-values are not represented.

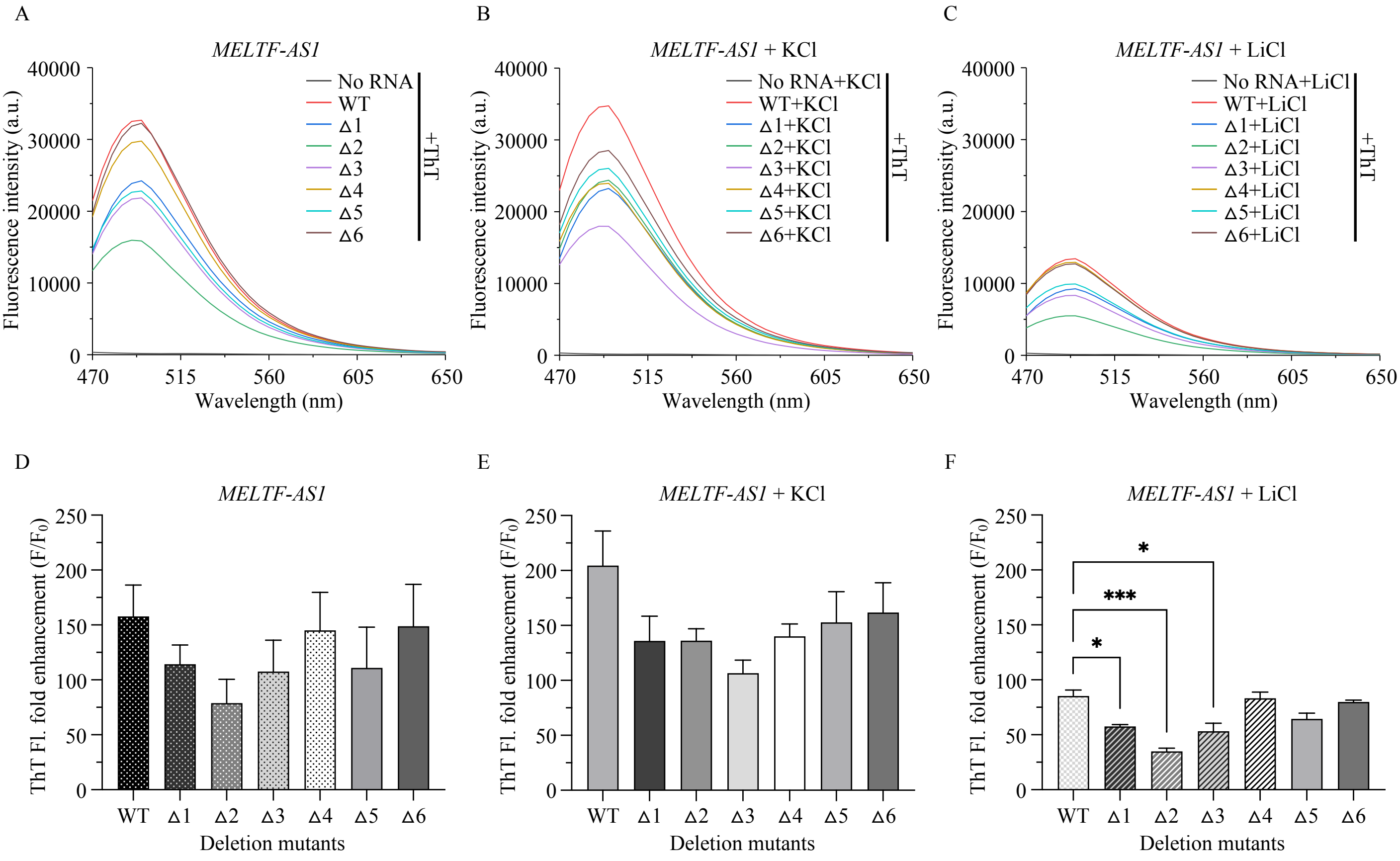


**Figure S7. Second, third, fourth, fifth, and sixth G-tracts are crucial for the stability of *MELTF-AS1* lncRNA G4.** A-F) ThT fluorescence enhancement assay of wild-type (WT) and deletion mutants (Δ) of RNAs (2 μM) folded in the presence or absence of 100 mM KCl or LiCl with ThT (2 μM). A-C) Increased mean fluorescence emission spectra, and D-F) fold enhancements in mean ± SEM ThT fluorescence at 488 nm when excited with 445 nm correspond to G4-formation in the presence or absence of specific monovalent cations ($K^+$ or $Li^+$). *P*-values: 0.0332 (*), 0.0021 (**), 0.0002 (***), <0.0001 (****). Non-significant P-values are not represented.

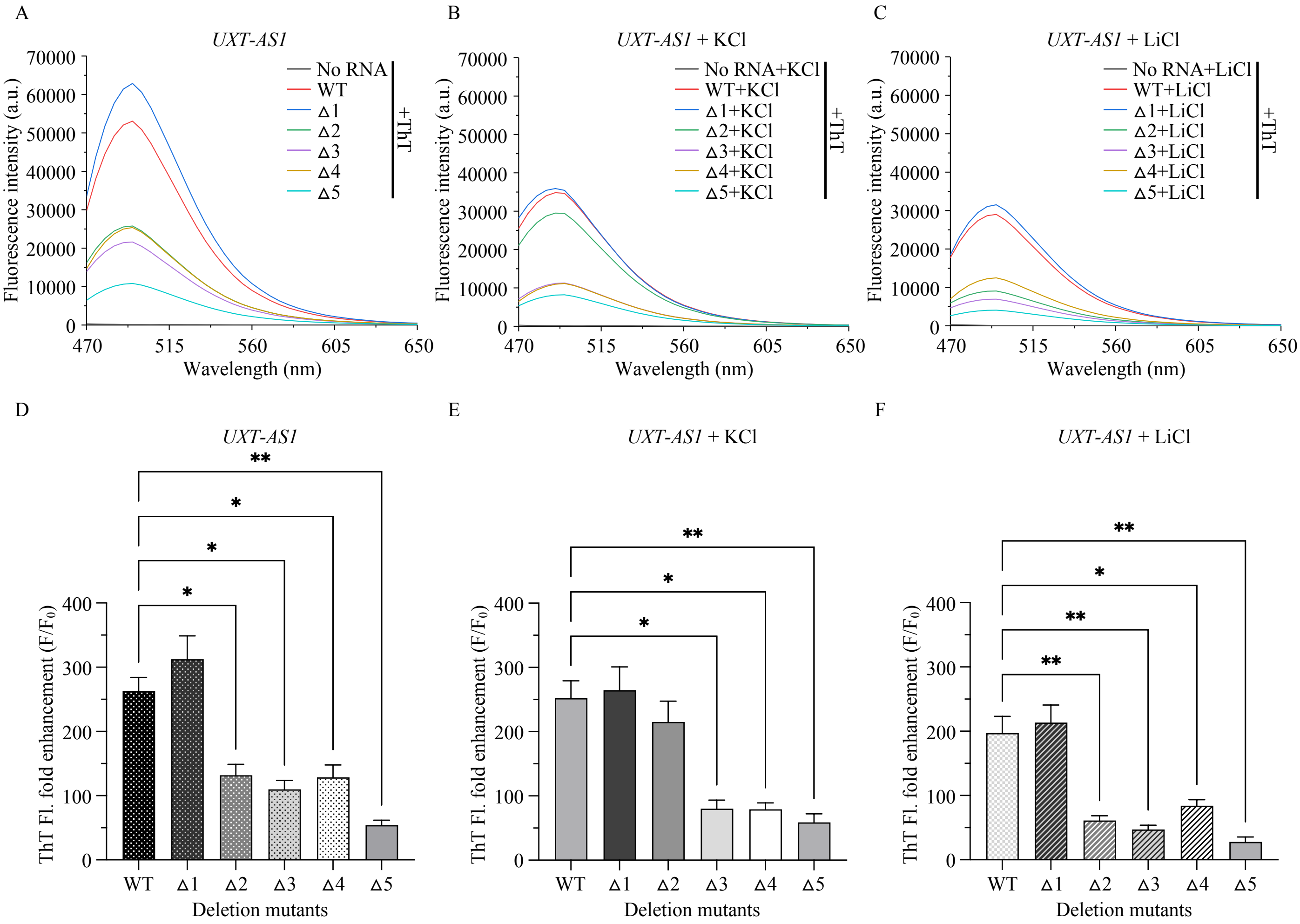


**Figure S8. Second, third, fourth, and fifth G-tracts are crucial for the stability of *UXT-AS1* lncRNA G4.** A-F) ThT fluorescence enhancement assay of wild-type (WT) and deletion mutants (Δ) of RNAs (2 μM) folded in the presence or absence of 100 mM KCl or LiCl with ThT (2 μM). A-C) Increased mean fluorescence emission spectra, and D-F) fold enhancements in mean ± SEM ThT fluorescence at 488 nm when excited with 445 nm correspond to G4-formation in the presence or absence of specific monovalent cations ($K^+$ or $Li^+$). *P*-values: 0.0332 (*), 0.0021 (**), 0.0002 (***), <0.0001 (****). Non-significant P-values are not represented.

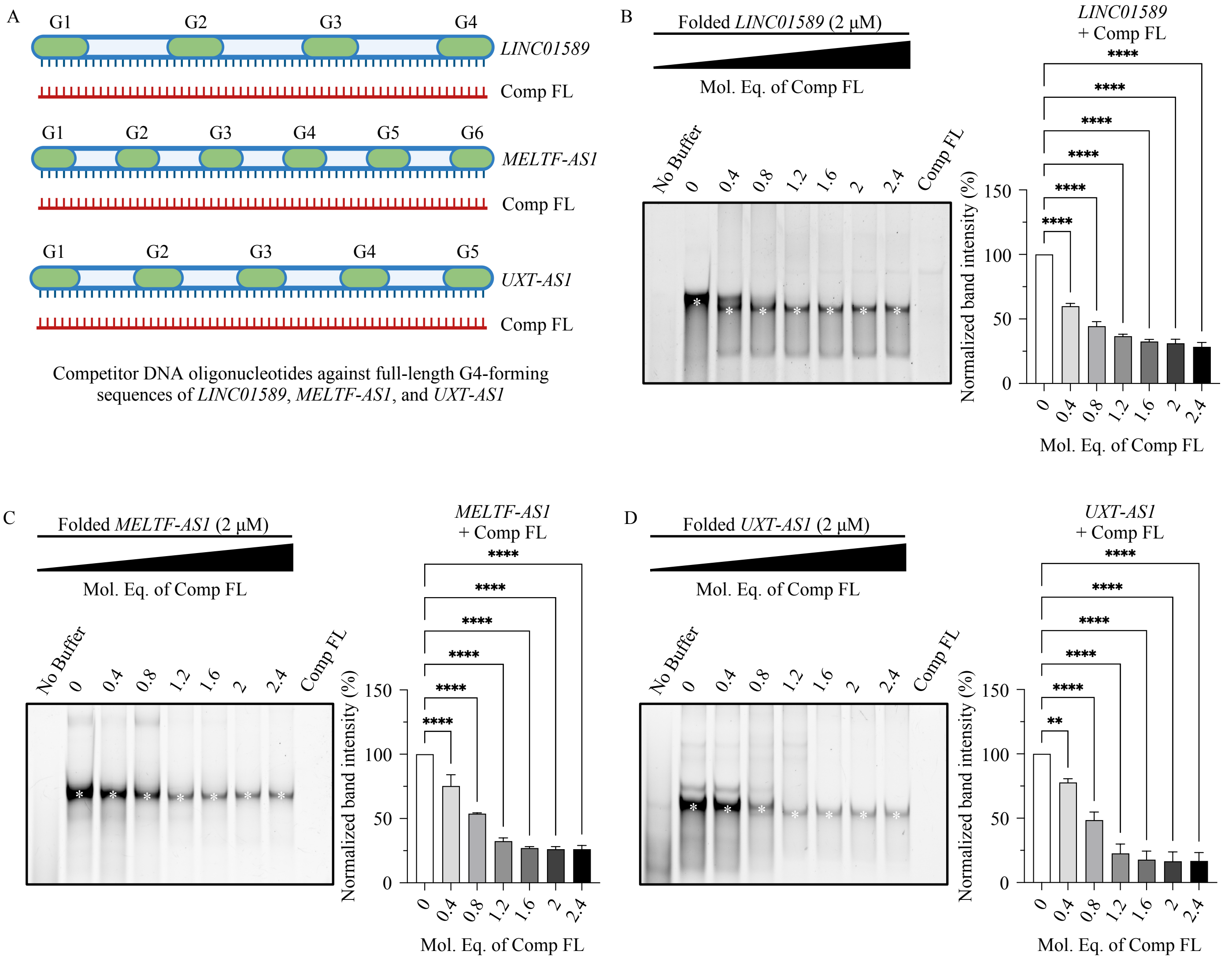


**Figure S9. Competitor Oligonucleotide Mediated-G4 Disruption (COM-G4D) indicating the loss of *LINC01589*, *MELTF-AS1*, and *UXT-AS1* lncRNA G4s.** A) COM-G4D using the competitor DNA oligonucleotides complementary to full-length G4-forming sequences of RNAs, binding to the cognate sequences, and blocking the participation of all G-tracts in G4-formation. B-D) Native PAGE (15%) of IVT RNAs (2 μM) folded in the presence of competitor DNA oligonucleotides (0 - 2.4 mol. eq.) and stained with ThT (0.5 μM) show bands (*) corresponding to G4s. Mean ± SD of their normalized intensities in the presence of competitors indicates G4-formation. *P*-values: $P \leq 0.05$, $P \leq 0.01$, $P \leq 0.001$, and $P \leq 0.0001$ are denoted with one asterisk (*), two asterisks (**), three asterisks (***), and four asterisks (****), respectively. Non-significant *P*-values are not represented.

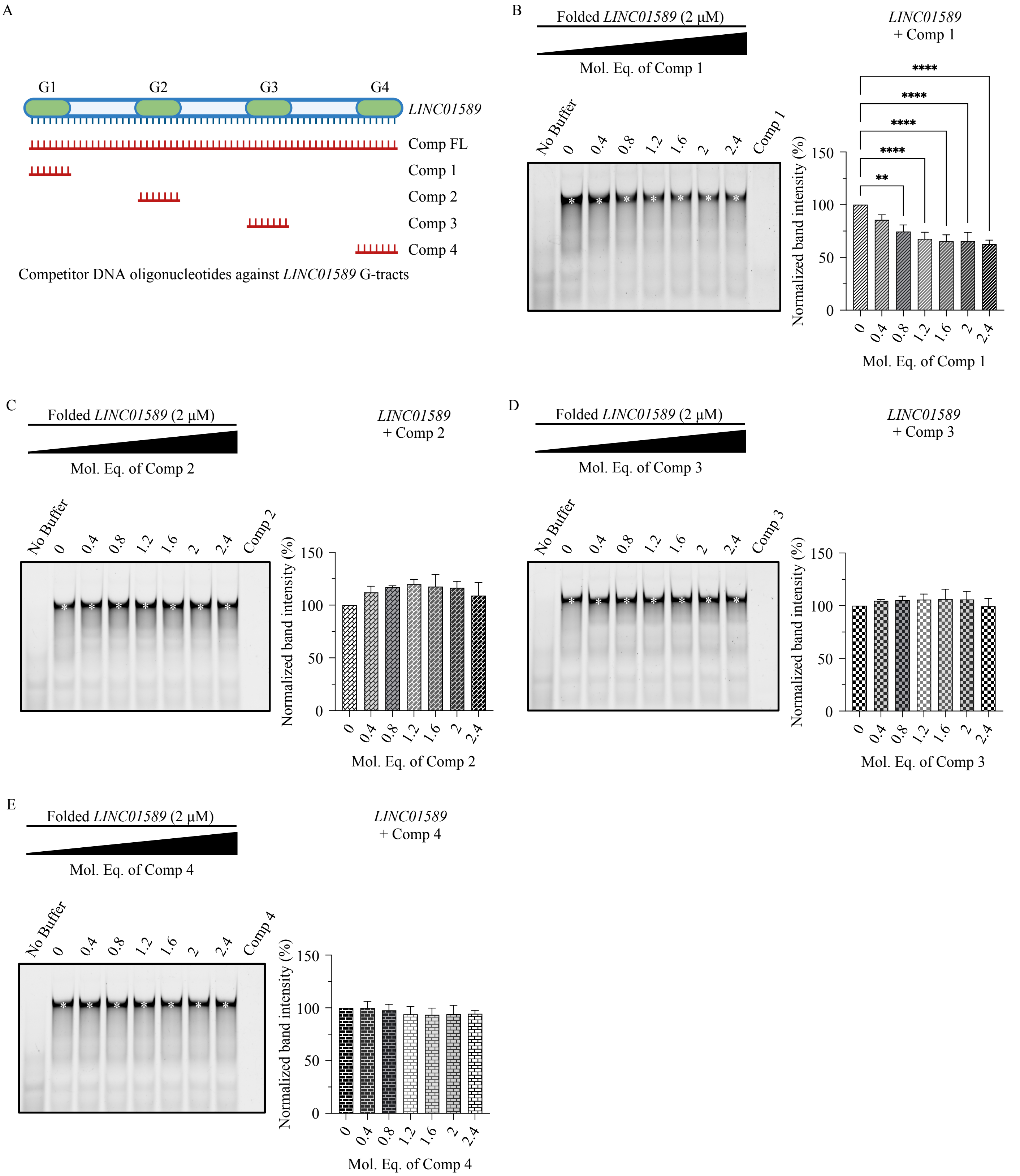


**Figure S10. COM-G4D on the G-tracts of *LINC01589* lncRNA G4.** A) COM-G4D using the competitor DNA oligonucleotides complementary to different G-tracts of RNA, binding to the cognate G-tract(s), and blocking their participation in G4-formation. B-E) Native PAGE (15%) of IVT RNAs (2 μM) folded in the presence of competitor DNA oligonucleotides (0 - 2.4 mol. eq.) and stained with ThT (0.5 μM) show bands (*) corresponding to G4s. Mean ± SD of their normalized intensities in the presence of each competitor indicates G4-formation. *P*-values: $P \leq 0.05$, $P \leq 0.01$, $P \leq 0.001$, and $P \leq 0.0001$ are denoted with one asterisk (*), two asterisks (**), three asterisks (***), and four asterisks (****), respectively. Non-significant *P*-values are not represented.

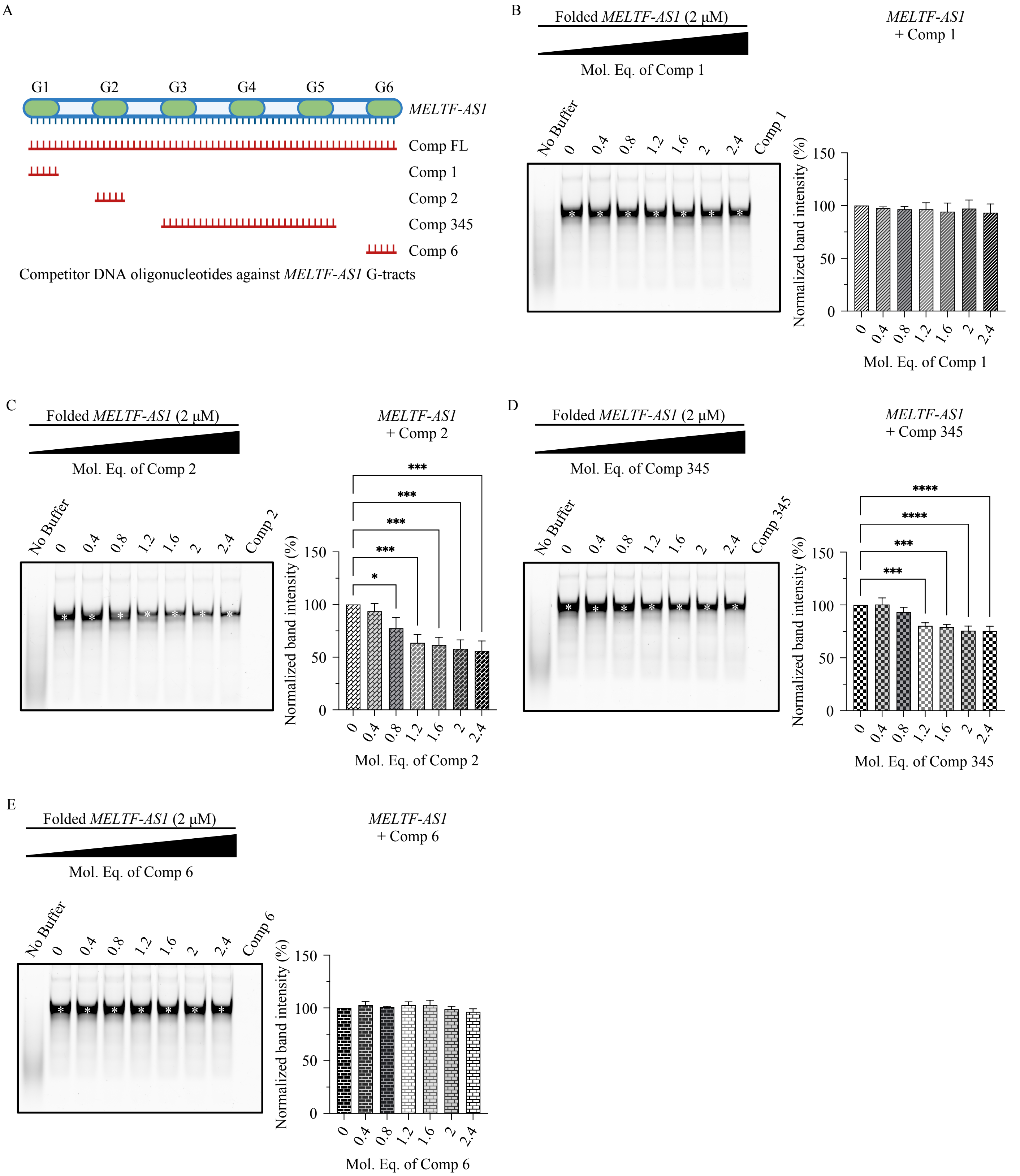


**Figure S11. COM-G4D on the G-tracts of *MELTF-AS1* lncRNA G4.** A) COM-G4D using the competitor DNA oligonucleotides complementary to different G-tracts of RNA, binding to the cognate G-tract(s), and blocking their participation in G4-formation. B-E) Native PAGE (15%) of IVT RNAs (2 μM) folded in the presence of competitor DNA oligonucleotides (0 - 2.4 mol. eq.) and stained with ThT (0.5 μM) show bands (*) corresponding to G4s. Mean ± SD of their normalized intensities in the presence of each competitor indicates G4-formation. *P*-values: $P \leq 0.05$, $P \leq 0.01$, $P \leq 0.001$, and $P \leq 0.0001$ are denoted with one asterisk (*), two asterisks (**), three asterisks (***), and four asterisks (****), respectively. Non-significant *P*-values are not represented.

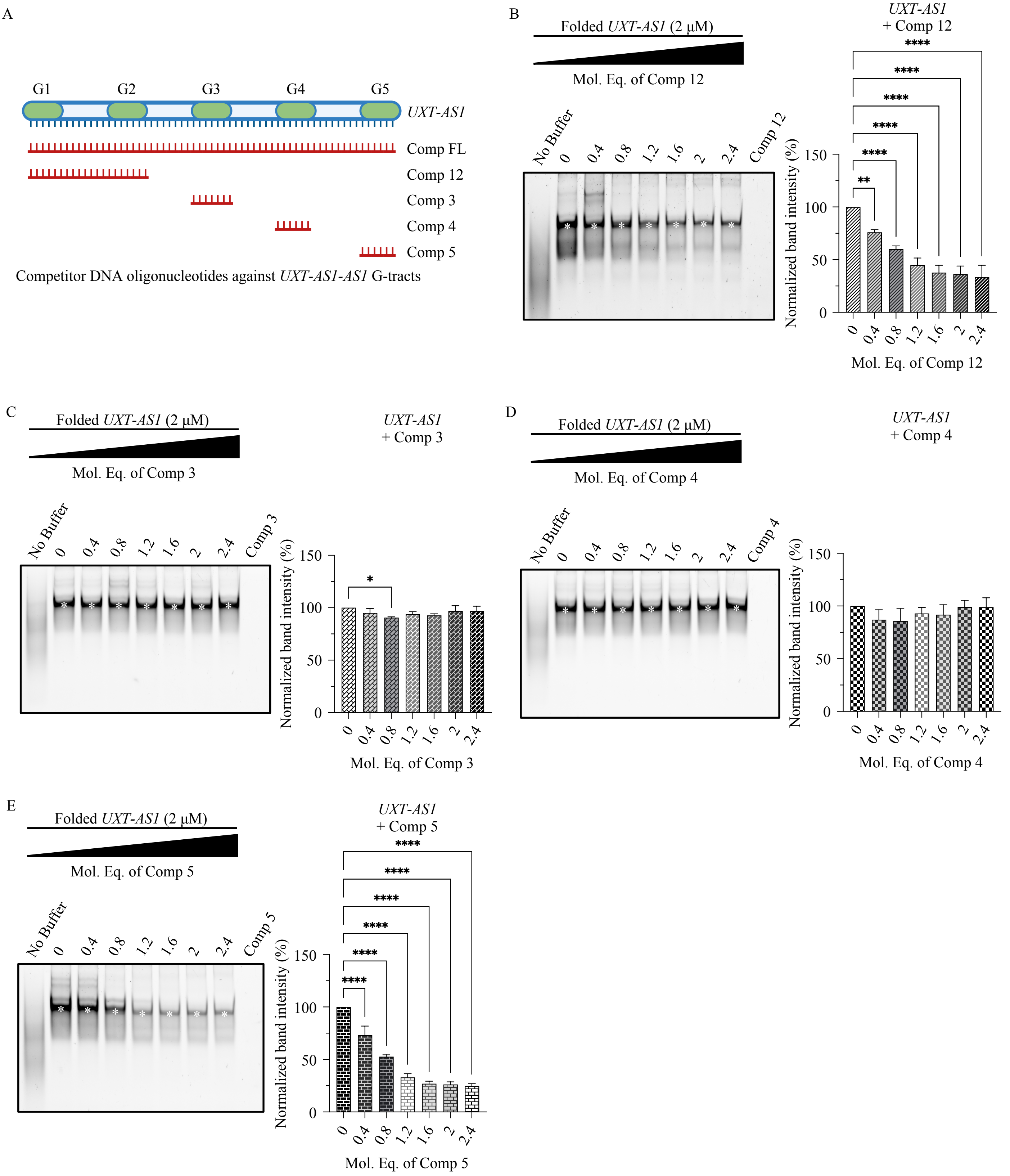


**Figure S12. COM-G4D on the G-tracts of *UXT-AS1* lncRNA G4.** A) COM-G4D using the competitor DNA oligonucleotides complementary to different G-tracts of RNA, binding to the cognate G-tract(s), and blocking their participation in G4-formation. B-E) Native PAGE (15%) of IVT RNAs (2 µM) folded in the presence of competitor DNA oligonucleotides (0 - 2.4 mol. eq.) and stained with ThT (0.5 µM) show bands (*) corresponding to G4s. Mean ± SD of their normalized intensities in the presence of each competitor indicates G4-formation. *P*-values: $P \leq 0.05$, $P \leq 0.01$, $P \leq 0.001$, and $P \leq 0.0001$ are denoted with one asterisk (*), two asterisks (**), three asterisks (***), and four asterisks (****), respectively. Non-significant *P*-values are not represented.

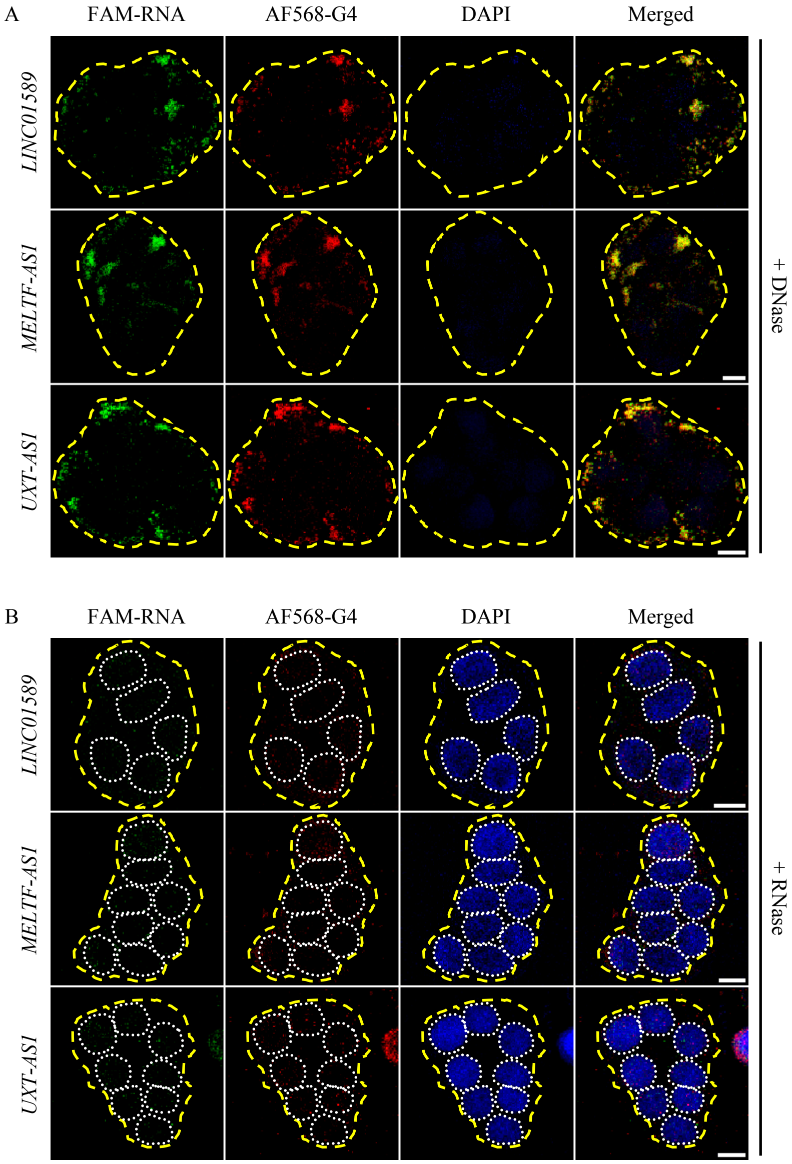


**Figure S13. RNA G4-Immuno-FISH identifies the G4s in *LINC01589*, *MELTF-AS1*, and *UXT-AS1* lncRNAs in CRC.** A-B) RNA G4-Immuno-FISH in HT-29 cells treated with A) DNase, and B) RNase, probed with G4-specific FLAG-BG4 – anti-FLAG – Alexa Fluor 568™ (AF568) antibodies, and hybridized with 5′ 6-FAM-labelled DNA probes. Cell clusters (marked with dashed yellow lines) imaged using Confocal microscopy (scale bar: 10 µm) in the FAM-RNA and AF568-G4 panels show the binding of 6-FAM-labelled DNA probes to the RNAs rather than DNAs, and the presence of G4s, respectively. Merged panel shows foci corresponding to RNAs and G4s, and colocalized foci corresponding to RNA G4s. DAPI staining indicates the nucleus (marked with white dotted lines).

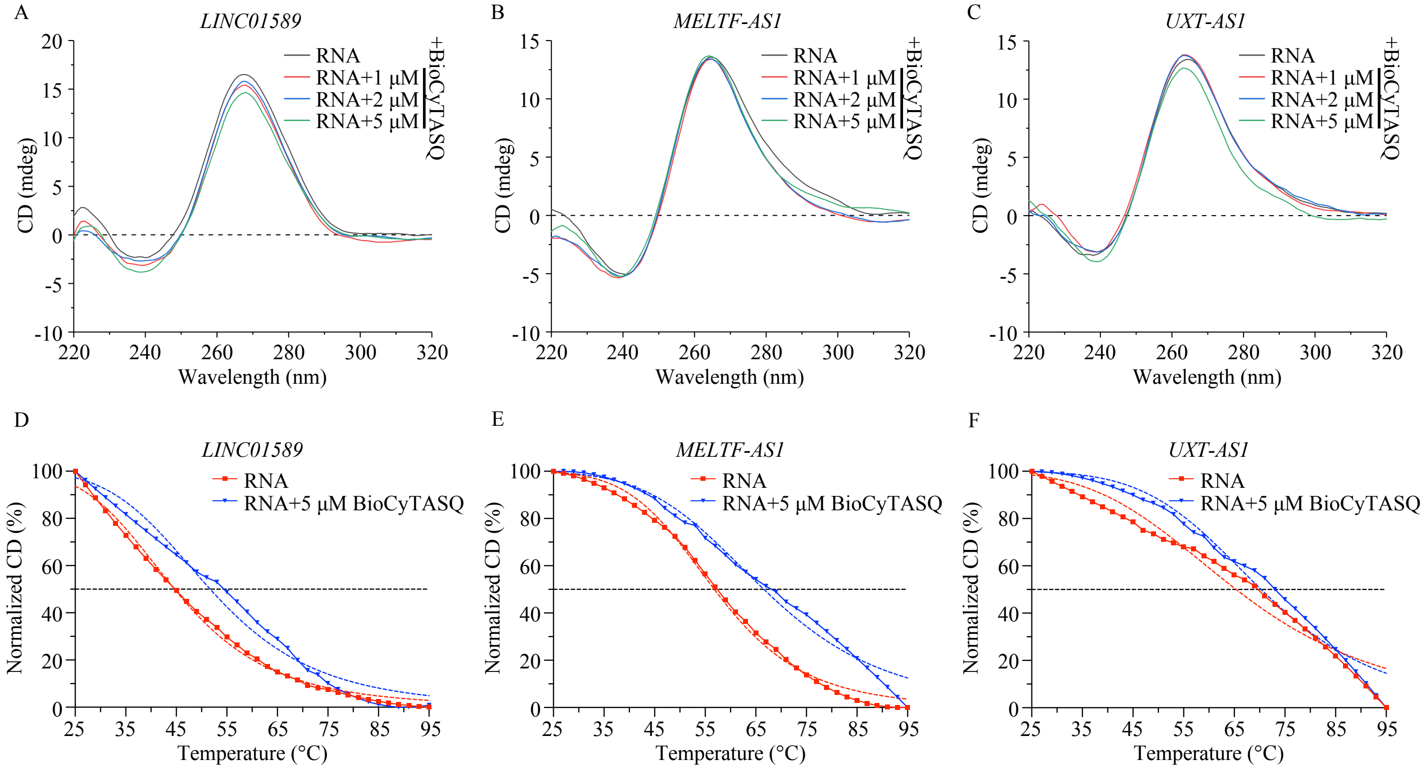


**Figure S14. BioCyTASQ used in G4RP-RT-qPCR stabilizes the *LINC01589*, *MELTF-AS1*, and *UXT-AS1* lncRNA G4s *in vitro*.** A-C) CD spectroscopy of folded synthetic RNAs (1 μM) titrated with BioCyTASQ (1 – 5 μM). No change in mean CD spectra corresponds to no effect of BioCyTASQ on the parallel G4 topologies. D-F) Normalized mean CD spectra of folded synthetic RNAs (1 μM) in the presence of BioCyTASQ (5 μM) at maxima obtained in the CD spectra, with increasing temperature. Greater thermal stability of RNAs in the presence of BioCyTASQ shows its stabilizing effect on the G4s. Dashed lines represent the plots fitted using the dose response-inhibition model of non-linear regression. *P*-values: $P \leq 0.05$, $P \leq 0.01$, $P \leq 0.001$, and $P \leq 0.0001$ are denoted with one asterisk (*), two asterisks (**), three asterisks (***), and four asterisks (****), respectively. Non-significant *P*-values are not represented.
