## Supplementary word and excel for "G-quadruplex formation in long non-coding RNAs dysregulated in colorectal cancer": SI text_LncRNA G4-CRC manuscript_Submitted to bioRxiv_05.07.2024_SS.pdf

### Materials and Methods

#### Thioflavin T (ThT) Fluorescence enhancement assay

Wild-type (WT) and deletion mutants (△) of RNAs (2 μM) folded in the presence or absence of an additional 100 mM KCl or LiCl were mixed with ThT (2 μM). ThT fluorescence emission spectra from 470 nm to 650 nm, along with endpoint fluorescence emission at 488 nm, were recorded with excitation at 445 nm, using a BioTek Cytation Hybrid Multimode Reader (Agilent Technologies International Pvt. Ltd., Manesar, India).[1] The data were recorded in triplicates across two independent studies. The mean fluorescence intensities (arbitrary units) of emission spectra were plotted against the wavelength after smoothening the data with the Savitzky-Golay method and 20 points of window using Origin (Pro). The mean ThT fluorescence fold-enhancements were plotted with standard error of mean against different monovalent cations used for the wild-type and deletion mutants of respective RNAs using GraphPad Software. Ordinary one-way or two-way ANOVA were employed for the statistical analyses.

#### Reverse Transcriptase stop (RT stop) assay

A 5′ Texas Red (Tx Red)-labelled primer was designed to bind to the primer binding site of IVT wild-type RNAs. To achieve separation between the primer binding site and the PQS, ten extra bases were included. Wild-type RNAs (2 μM) were folded in an annealing buffer: 10 mM Tris-Cl (pH 7.5), 50 mM NaCl and 1 mM EDTA (pH 8.0), in the presence or absence of additional 50, 100, and 150 mM KCl or LiCl, along with dNTPs (2 mM), and a 5′ Tx Red-labelled primer (100 nM). The reverse transcription of the folded RNAs into the cDNAs was performed in an extension buffer: 10 mM Tris-Cl (pH 7.5) and 0.1 mM EDTA (pH 8.0), supplemented with 3 mM $MgCl_2$ and M-MLV Reverse Transcriptase (RNase H Minus) (RT) (4U/ μl) at 37 °C for 60 minutes. The reaction was terminated by mixing the reverse transcribed products with 2X denaturing loading dye to a working concentration of 1X: 47.5% formamide, 5 mM EDTA (pH 8.0), 0.05% Xylene Cyanol and 0.05 % Bromophenol Blue, and incubating the samples at 95 °C for 10 minutes. The resulting samples were run on a 15% denaturing PAGE (8 M urea).[2–6] Gel imaging was performed using the ChemiDoc™ MP Imaging System under Green Epi at 602/50 nm, with a 10-minute exposure. The data were recorded across three independent studies. The intensities of the full-length cDNA bands were quantified using ImageJ.[7] The means of normalized band intensities were plotted with standard deviation against varying concentrations of monovalent cations used for the respective RNAs using GraphPad Software. Ordinary one-way ANOVA was applied for the statistical analyses.

**Native Polyacrylamide Gel Electrophoresis of deletion mutants of RNAs**

Folded wild-type and deletion mutants of RNAs (2 μM) were mixed with 6X Gel loading dye (Catalog no. ML015, HiMedia Laboratories Pvt. Ltd., Thane-West, India) to a working concentration of 1X. The 15% native polyacrylamide gel was prepared using 40% Acrylamide/Bisacrylamide Solution (Catalog no. ML083, HiMedia Laboratories Pvt. Ltd., Thane-West, India) in 1X Tris-borate-EDTA (TBE) buffer. The RNAs, mixed with dye, were loaded in the gel, and electrophoresis was conducted at 150 V for 60 minutes in 1X TBE buffer. Subsequently, the gel was stained with Thioflavin T (ThT) (0.5 μM) in 1X TBE buffer for 15 minutes. Gel imaging was performed using the

ChemiDoc™ MP Imaging System (Bio-Rad Laboratories India Pvt. Ltd., Gurugram, India) at 532/28 nm, Blue Epi, with a 30-second exposure. The data were recorded across three independent studies. The intensities of the bands were quantified using ImageJ. The means of normalized band intensities were plotted with standard deviation against the wild-type and deletion mutants of respective RNAs using GraphPad Software. Ordinary one-way ANOVA was applied for the statistical analyses.

**Competitor Oligonucleotide Mediated-G4 Disruption (COM-G4D)**

Competitor DNA oligonucleotides were designed to be complementary to the full-length G4-forming sequences or different G-tracts of the wild-type RNAs, as listed in Table S3. Due to the very short flanking regions of the third, fourth, and fifth G-tracts of *MELTF-AS1* lncRNA and, first and second G-tracts of *UXT-AS1* lncRNA, the design of independent competitor DNAs complementary to each of these G-tracts was not ideal. Hence, a single competitor DNAs complementary to these cognate G-tracts (*MELTF-AS1*_Comp 345 and *UXT-AS1*_Comp 12) were used to assess their consolidated effect. Wild-type RNAs (2 μM) were folded in the presence of increasing concentrations (0 - 2.4 mol. equiv) of competitor DNA oligonucleotides. Folded RNAs were mixed with 6X Gel loading dye to a working concentration of 1X, and run on a 15% native PAGE. The ThT was used for gel staining, and gel imaging was carried out using the ChemiDoc™ MP Imaging System, as described earlier. The data were recorded across three independent studies. The intensities of the bands were quantified using ImageJ. The means of normalized band intensities were plotted with standard deviation against varying concentrations of competitor DNA oligonucleotides used for the respective RNAs using GraphPad Software. Ordinary one-way ANOVA was applied for the statistical analyses.

**Statistical significance**

The statistical analyses included appropriate tests, and the resulting statistical significance ($P$-values) with $P \leq 0.05$, $P \leq 0.01$, $P \leq 0.001$, and $P \leq 0.0001$ are denoted with one asterisk (*), two asterisks

(**), three asterisks (***), and four asterisks (****), respectively. Non-significant *P*-values are not represented.

### Results and Discussion

#### ***LINC01589*, *MELTF-AS1*, and *UXT-AS1* lncRNA G4s stability is affected by specific monovalent cations**

We investigated the influence of specific monovalent cations on the G4s formed by *LINC01589*, *MELTF-AS1*, and *UXT-AS1* lncRNAs *via* Thioflavin T (ThT) fluorescence enhancement assay (Figure S3A). An enhancement of 90-, 158-, and 263-fold in ThT fluorescence was observed for the *LINC01589*, *MELTF-AS1*, and *UXT-AS1* lncRNAs, respectively, without supplementation of KCl or LiCl (Figure S3B-G).[1] Similar to the CD-melting behaviour, the pattern of fluorescence enhancement for *LINC01589*, *MELTF-AS1*, and *UXT-AS1* lncRNAs is consistent with the superlative stability potentially associated with a higher number of G-quartets (2G, 3G, and 4G, respectively).[8,9] We obtained surprising results when assessing the influence of specific monovalent cations ($K^+$ and $Li^+$) on the stability of G4s formed by these lncRNAs (Figure S3B-G). While the supplementation of LiCl exhibited an apparent stabilizing effect than KCl for *LINC01589*, KCl supplementation demonstrated noticeable stabilization for *MELTF-AS1* compared to LiCl (Figure S3B, C, E, F). Conversely, addition of neither KCl nor LiCl had any observable effect on *UXT-AS1* (Figure S3D, G). Although there is a popular notion that $K^+$ stabilizes G4s and $Li^+$ destabilizes them, a few reports suggest the destabilizing effect of $K^+$ and the neutral effect of $Li^+$ on specific G4s, highlighting the varied effects of these monovalent cations on G4s.[10–12] Notably, the possible existence of the traces of monovalent cations in the folding buffer without supplementation of KCl or LiCl might be sufficient to promote the spontaneous folding of these G4s in a stable manner. Consequently, the supplementation of KCl or LiCl did not result in a logarithmic fold-change in G4-stability.

For further investigation, we conducted a modified Reverse Transcriptase stop (RT stop) assay by utilizing a primer binding sequence at the 3′-end of IVT-derived RNAs that can hybridize with a 5′ Texas Red (Tx Red)-labelled primer to label the reverse transcribed cDNAs (Figure S4A; Table S3).[2–6] Since this *in vitro* investigation aims to identify the fundamental cationic contributor of G4-stability, mimicking the complex physiological cationic environment was not intended in our experimental setup. Hence, the experiments were performed in the presence or absence of an additional 50, 100, or 150 mM KCl or LiCl. The $K^+$ and $Li^+$ ions influenced the reverse transcription of all lncRNA G4s into full-length cDNAs, exhibiting inconsistent effects (Figure S4B-G). Notably, the varied cationic concentrations and the type of Reverse Transcriptase (RT) employed might affect the processivity of enzyme and the experimental resolution, respectively. Given the sensitivity of G4 folding and stability to the ionic composition of buffers, the observed effects of K+ and Li+ collectively indicate a modest, concentration-dependent modulation of G4s formed by these lncRNAs, warranting further investigation.

**Influence of G-tracts on the stability of *LINC01589*, *MELTF-AS1*, and *UXT-AS1* lncRNA G4s**

We next utilized the IVT-derived wild-type (WT) lncRNAs and singular G-tract devoid deletion mutants (Δ) of lncRNAs to examine the contribution of individual G-tracts within these lncRNAs in maintaining the G4-stability. For this, we analyzed them on 15% native PAGE stained with ThT (Figure S5A-C; Table S3).[13] The significant contribution of the first G-tract of *LINC01589* lncRNA towards maintaining the G4-stability was suggested by a faint ThT-stained band observed only in the Δ1 compared to the wild-type lncRNA (Figure S5A, D). Notably, the presence of four guanines in the third G-tract of *LINC01589* can potentially provide two independent G-tracts with two guanines each and no loop (Figure S5A). One of these two G-tracts can act as a "spare tire" by substituting when either the first, second, or fourth G-tract of *LINC01589* is deleted, thereby fulfilling the requirement of four G-tracts for the G4-formation.[14] Furthermore, the formation of bulges can lead to G4 topologies from non-conventional motifs by bulging out intervening nucleotide(s), thereby connecting adjacent guanines in the same column or strand of the G-quartet core. This bulging results in the tandem arrangement of non-continuous or distant guanines with intervening nucleotide(s) into a new G-tract, forming a continuous column that supports the G-quartet core and thus facilitates G4-formation.[15,16] Similarly, when the third G-tract of *LINC01589* containing four guanines is deleted, the intervening nucleotide(s) between two adjacent guanines can potentially bulge out to form a new G-tract, which can then act as a "spare tire" to replace the deleted third G-tract (Figure S5A). *MELTF-AS1* highlighted the critical contribution of its second, third, fourth, fifth, and sixth G-tracts towards stable G4-folding as the band intensities for all deletion mutants of this lncRNA, except Δ1, decreased significantly compared to the wild-type lncRNA (Figure S5B, E). For *UXT-AS1* lncRNA, an important contribution of the second, third, fourth, and fifth G-tracts in maintaining the stability of G4 was pointed by faint ThT-stained bands for all deletion mutants, except Δ1, compared to the wild-type lncRNA (Figure S5C, F). The deletion mutants of lncRNAs also indicated their reduced ability to form G4s at levels similar to wild-type lncRNA in the ThT fluorescence enhancement assay (Figure S6-8). These findings underscore the significance of first G-tract of *LINC01589*, and all except the first G-tract of *MELTF-AS1* and *UXT-AS1* lncRNAs towards maintaining the stability of cognate G4s.

A novel technique termed Competitor Oligonucleotide Mediated-G4 Disruption (COM-G4D) was used to further explore the G4-formation within lncRNAs. It is an extended application of antisense oligonucleotide hybridization, which involves the use of increasing concentrations of competitor DNA oligonucleotides complementary to the full-length G4-forming sequences or different G-tract(s) of lncRNA G4s. These competitor DNAs can bind and block the participation of all or selective G-tracts in G4-formation, thereby influencing G4-formation in each case (Figure S9A; Table S3). The samples were run on 15% native PAGE stained with ThT. The increasing concentrations of full-length competitor DNAs for *LINC01589*, *MELTF-AS1*, and *UXT-AS1* lncRNAs (*LINC01589*_Comp FL, *MELTF-AS1*_Comp FL, and *UXT-AS1*_Comp FL, respectively) led to a significant reduction in the band intensities of cognate lncRNA G4s compared to those without competitors (Figure S9B-D).

This finding unequivocally demonstrates the capacity of complementary DNA oligonucleotides to influence the G4-forming capability of these lncRNAs.

Based on the observed behaviour of deletion mutants and COM-G4D in the presence of full-length competitor DNAs, we attempted to examine the performance of short competitor DNAs complementary to individual G-tracts of each lncRNA (Figure S10A, S11A, S12A; Table S3). Decreased band intensities were observed with the increasing concentrations of *LINC01589*_Comp 1, *MELTF-AS1*_Comp 2, *UXT-AS1*_Comp 12, and *UXT-AS1*_Comp 5 (Figure S10B, S11C, S12B, E). This observation suggests that these specific competitor DNAs bind to and disrupt the respective lncRNA G4s, highlighting the critical role of these G-tracts in maintaining G4-stability. In contrast, the other competitor DNAs did not produce similar changes in band intensities (Figure S10-12). The use of competitor DNAs complementary to specific G-tracts could result in the following outcomes: (i) binding leading to G4 disruption, (ii) binding without G4 disruption, and (iii) inability to bind, possibly due to the small length of competitor DNAs, and no G4 disruption. While outcome (i) could inform about the contribution of specific G-tracts towards G4-formation, outcomes (ii) and (iii) could create ambiguities in interpretation. Nevertheless, the outcomes observed with *LINC01589*_Comp 1, *MELTF-AS1*_Comp 2, *UXT-AS1*_Comp 12, and *UXT-AS1*_Comp 5 are consistent with those seen in corresponding deletion mutants. With further exploratory investigations, COM-G4D has the potential to be used as a tool to determine the importance of individual G-tracts in G4-formation.
